# Dystonia-associated TorsinA-ΔE mutation induces a gain-of-function interaction with XPO1 via its N-Terminal hydrophobic segment

**DOI:** 10.64898/2026.08.17.745292

**Authors:** Haochen Cui, Yuntian Duan, Md Kobirul Islam, Md Abir Hosain, Jiyu Li, Xiaohong Lu, Baojin Ding

**Affiliations:** Department of Biochemistry and Molecular Biology, Louisiana State University Health Sciences Center at Shreveport, Shreveport, LA 71103, USA; Department of Pharmacology, Toxicology, and Neuroscience, Louisiana State University Health Sciences Center at Shreveport, Shreveport, LA 71103, USA

**Keywords:** TorsinA, DYT1 dystonia, exportin 1, nucleocytoplasmic transport, proteomics, motor neurons, AlphaFold, peptides

## Abstract

Childhood-onset DYT1 dystonia is a neurodevelopmental movement disorder caused by a three-base-pair deletion (ΔGAG; ΔE) in the *TOR1A* gene, which encodes TorsinA, a membrane-associated AAA+ (ATPase associated with diverse cellular activities) ATPase. However, the mechanisms by which the ΔE mutation causes neuronal dysfunction remain poorly understood. Using patient-derived neurons, we previously demonstrated that TorsinA-ΔE disrupts the nucleocytoplasmic transport (NCT) of both RNA and protein cargos. In the present study, proteomic analysis of induced human motor neurons revealed a markedly enhanced association between ΔE and exportin 1 (XPO1), a major nuclear export receptor. This aberrant association was enriched at the nuclear envelope and accompanied by impaired XPO1-mediated nuclear export. By integrating AlphaFold-based structural modeling with molecular, biochemical, and cellular analyses, we identified the N-terminal hydrophobic segment (HS) of TorsinA as a critical contributor to its interaction with XPO1. Deletion of the HS from ΔE reduced its association with XPO1, altered its nuclear envelope enrichment, and restored nuclear export. Moreover, expression of HS-derived peptides in patient-derived DYT1 neurons improved nuclear export, neurite outgrowth and branching, maturation-associated gene expression, and neuronal survival. Together, these findings identify an aberrant gain-of-function association between TorsinA-ΔE and XPO1 as a mechanism contributing to NCT dysfunction in DYT1 dystonia and establish the HS-dependent ΔE-XPO1 interaction as a potential therapeutic target.

**Significance Statement:** DYT1 dystonia is a childhood-onset movement disorder caused by a mutation in TorsinA, but how this mutation disrupts neuronal function remains unclear. We identify an abnormally enhanced interaction between mutant TorsinA and exportin 1 (XPO1), a protein that transports cargo from the nucleus. This interaction depends on the N-terminal hydrophobic segment of TorsinA and contributes to defective nuclear transport in DYT1 neurons. Deleting this segment or expressing short peptides derived from it improved nuclear transport, neurite growth, neuronal maturation, and survival in patient-derived neurons. These findings reveal a gain-of-function mechanism that complements the established loss-of-function model of DYT1 dystonia and identify the mutant TorsinA-XPO1 interaction as a potential therapeutic target.

## Introduction

Childhood-onset DYT1 dystonia is the most common forms of hereditary isolated dystonia and is often associated with severe motor impairment (1, 2). Symptoms typically emerge during childhood or adolescence, a critical period for motor development and learning (3–5). DYT1 dystonia is most commonly caused by a heterozygous three-base-pair deletion (ΔGAG) in exon 5 of *TOR1A*, resulting in the loss of a glutamate residue near the C terminus of TorsinA (ΔE) (2, 6, 7). Although ΔE has generally been considered a loss-of-function mutation (8–10), the molecular mechanisms through which it causes neuronal dysfunction remain poorly understood.

Torsin proteins belong to the conserved AAA+ (ATPases Associated with diverse cellular Activities) superfamily, whose members harness ATP hydrolysis to remodel or disassemble protein complexes involved in processes such as membrane trafficking, cytoskeletal dynamics, vesicle fusion, and stress responses (11–13). TorsinA shares homology with the bacterial Clp/Hsp100 family, canonical AAA+ ATPases (14, 15) but is considered noncanonical because of two distinctive properties. First, torsins are the only known AAA+ proteins localized within the endoplasmic reticulum (ER) and nuclear envelope (NE) (16, 17). Second, they lack intrinsic ATPase activity and require the ER/NE transmembrane cofactors LAP1 and LULL1 (also known as TOR1AIP1 and TOR1AIP2) for enzymatic activation (10, 18, 19). These features suggest that TorsinA primarily functions as part of a membrane-associated protein complex.

Nevertheless, several findings suggest that TorsinA may also act outside this canonical cofactor-dependent context. Oxidative and heat-shock stresses have been reported to induce the redistribution of TorsinA from the ER to the cytosol (20, 21). In addition, the C-terminus of TorsinA binds to cytosolic proteins, including JNK-interacting protein 1 (JIP1) and kinesin light chain 1 (KLC1) (22, 23). Recent studies using *Drosophila* indicated that dTor and the activator dLap1 differentially affect the fly fat body (24), and neither ATP binding nor hydrolysis by dTor is essential for germ cell development (25). These observations raise the possibility that a cytosolic pool of endogenous TorsinA exists and may perform functions independent of its canonical cofactors or ATPase activity.

Consistent with its unusual properties and subcellular localization, TorsinA has been implicated in diverse cellular processes, including nucleocytoskeletal coupling (26, 27), lipid metabolism (28–30), protein quality control (31–33), nuclear pore complex maturation and nucleocytoplasmic transport (NCT) (25, 34–36). Despite this expanding range of proposed functions, the precise molecular roles of TorsinA in cellular and neuronal physiology, particularly in human neurons, remain incompletely defined.

We previously modeled DYT1 dystonia using patient-derived neurons generated either through direct fibroblast-to-neuron conversion (37, 38) or through human induced pluripotent stem cell (hiPSC)-based differentiation (39–42). These neurons retain the endogenous ΔGAG mutation and recapitulate disease-relevant phenotypes, including nuclear deformation, Lamin B1 mislocalization, and impaired NCT (35, 36, 43). These observations suggest that disruption of NE organization and nuclear transport represents an important component of DYT1 pathogenesis. However, the molecular link between TorsinA-ΔE and the nuclear transport machinery remains unknown.

To define the mechanism underlying these abnormalities, we performed co-immunoprecipitation coupled with mass spectrometry (Co-IP/MS) in induced human motor neurons. This analysis identified a markedly enhanced association between ΔE and exportin 1 (XPO1), a major nuclear export receptor responsible for transporting nuclear export signal (NES)-containing protein cargos. By integrating AlphaFold-based structural modeling with molecular, biochemical, and cellular analyses, we identified the N-terminal hydrophobic segment (HS) of TorsinA as a critical contributor to its association with XPO1. Deletion of the HS from ΔE reduced its association with XPO1, altered its NE enrichment, and restored nuclear export. Moreover, expression of HS-derived peptides ameliorated nuclear export and neuronal deficits in patient-derived DYT1 neurons. Together, these findings support an aberrant gain-of-function mechanism in which enhanced HS-dependent association of ΔE with XPO1 contributes to NCT dysfunction. They establish a molecular link between the DYT1 mutation and the nuclear export machinery and identify the ΔE-XPO1 interaction as a potential therapeutic target.

## Results

### Proteomic analysis identifies altered ΔE-interacting proteins in induced human motor neurons

Using patient-derived neurons, we previously demonstrated that TorsinA-ΔE disrupts nuclear transport of both RNA and protein cargos (35). Notably, ΔE is highly enriched at the NE, whereas wild-type (WT) TorsinA is more broadly distributed in the cytoplasm (44–46). This distinct subcellular localization suggests that ΔE may engage in a different set of protein-protein interactions. Such altered interactions could contribute to the NCT defects and neuronal dysfunction observed in DYT1 dystonia.

To identify proteins that interact differentially with WT and mutant TorsinA-ΔE, we performed co-immunoprecipitation followed by mass spectrometry (Co-IP/MS) in human induced pluripotent stem cell (hiPSC)-derived motor neurons (MNs) (**Figure 1A**). Because endogenous TorsinA is expressed at low levels and suitable immunoprecipitation-grade antibodies are limited, we transduced neural progenitor cells with lentiviral vectors expressing GFP, WT TorsinA-GFP, or ΔE-GFP and subsequently differentiated them into MNs (**Figure S1**) (36, 40, 41). Consistent with previous reports (44, 45), ΔE-GFP accumulated at the NE, whereas WT TorsinA-GFP was broadly cytoplasmic (**Figure 1B**). To enhance immunoprecipitation efficiency and specificity, we used GFP-Trap, an anti-GFP nanobody conjugated to agarose beads. Coomassie Brilliant Blue staining of SDS-PAGE gels confirmed the expression of GFP, TorsinA-GFP, and ΔE-GFP at their expected molecular weights and their enrichment in the immunoprecipitated fractions (**Figure 1C**). Because overexpression of ΔE-GFP reduced neuronal survival and consequently lowered the immunoprecipitation yield, additional starting material was used for the ΔE-GFP condition. For mass-spectrometry analysis, protein abundance was normalized to the abundance of the corresponding GFP-tagged bait in each sample.

**Figure 1.**
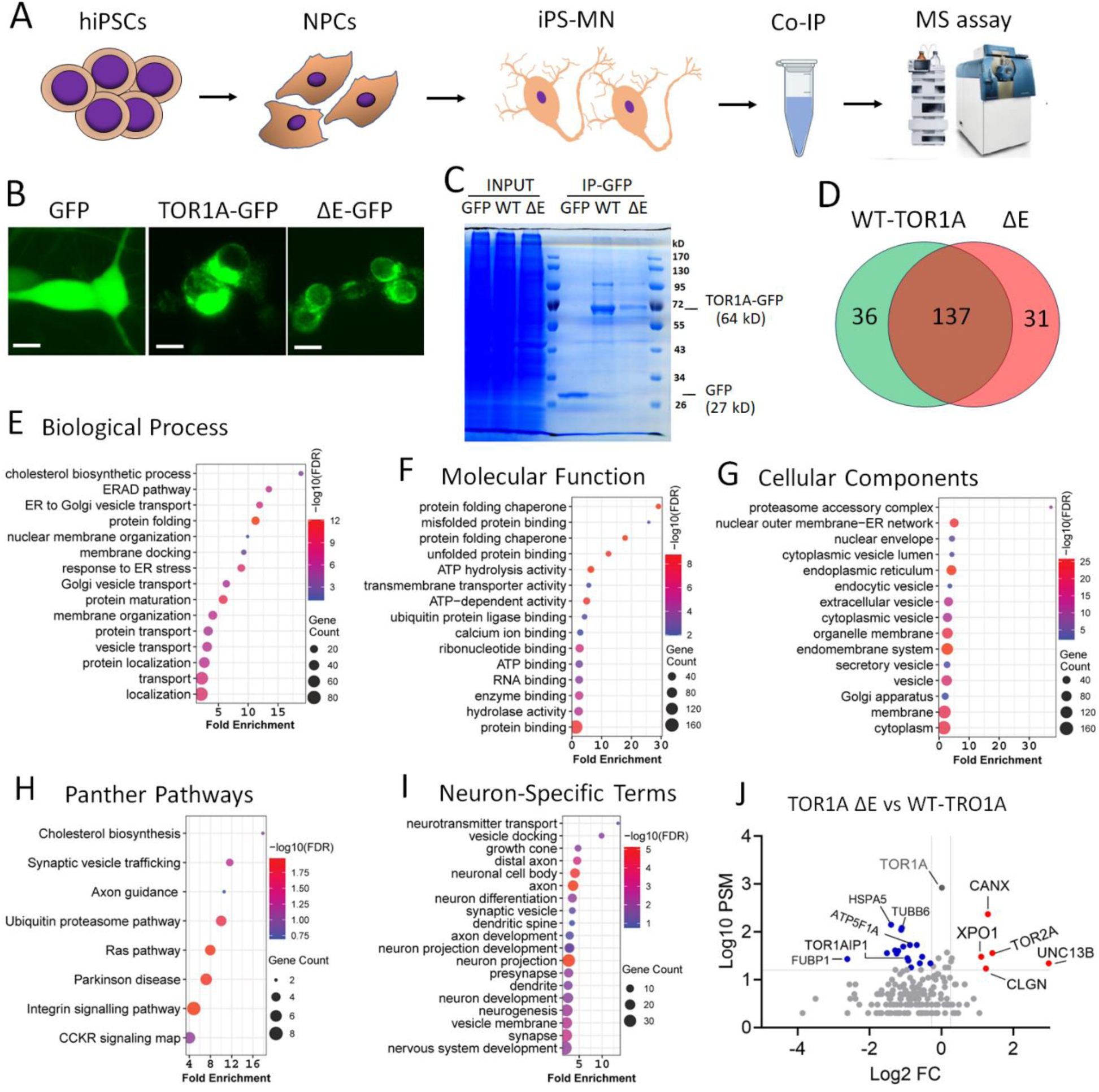
Identification of proteins differentially associated with torsinA-ΔE in induced human motor neurons. **(A)** Schematic illustrating the workflow for generating hiPSC-derived motor neurons (iMNs) and performing co-immunoprecipitation coupled with mass spectrometry (Co-IP/MS). **(B)** Representative fluorescence images of iMNs expressing GFP, WT TOR1A-GFP, or ΔE-GFP at 8 days post-viral induction (DPI). Scale bars, 10 µm. **(C)** Coomassie Brilliant Blue-stained SDS-PAGE gel showing input (1% of total lysis) and immunoprecipitated (IP; 5% of total eluate) fractions from iMNs expressing the indicated proteins. **(D)** Venn diagram showing proteins associated with WT, ΔE, or both, identified using a fivefold enrichment threshold relative to the GFP control. (**E–I**) Gene Ontology (GO) and pathway enrichment analyses of the 204 TorsinA-associated proteins identified by Co-IP/MS, including 36 detected only with WT TorsinA, 31 detected only with ΔE, and 137 detected with both: **(E)** biological processes, **(F)** molecular functions, **(G)** cellular components, **(H)** PANTHER pathways, and **(I)** neuron-related GO terms. (**J**) Volcano plot comparing the relative association of proteins with WT and ΔE in iMNs.

Using a fivefold enrichment threshold relative to the GFP control, we identified 204 TorsinA-associated proteins among 1,085 detected candidates. Of these, 36 were detected only with WT TorsinA, 31 only with ΔE, and 137 with both proteins (**Figure 1D** and **Table S6**). Gene Ontology (GO) analysis (**Table S7**) showed that these TorsinA-associated proteins were enriched in biological processes involving protein folding and maturation, intracellular trafficking, the ER stress response, ER-to-Golgi transport, ER-associated degradation, Golgi vesicle transport, cholesterol biosynthesis, and nuclear membrane organization (**Figure 1E**). Enriched molecular functions included ATP-dependent activity, protein-folding chaperone activity, enzyme binding, ubiquitin ligase binding, and transmembrane transporter activity (**Figure 1F**). Consistent with these functions, the interacting proteins were predominantly associated with the endomembrane system, ER, NE-ER network, vesicles, and vesicle lumen (**Figure 1G**). PANTHER pathway analysis further identified enrichment in the ubiquitin-proteasome system, Parkinson disease pathways, integrin and Ras signaling, synaptic vesicle trafficking, cholesterol biosynthesis, and axon guidance (**Figure 1H**).

These results are consistent with previous studies showing that TorsinA resides predominantly within the ER and NE and participates in membrane-cytoskeleton interactions, vesicle trafficking, protein quality control, and lipid metabolism (3, 18, 47–50). The identified proteins were also enriched in neuron-related biological processes, including neurogenesis, neuronal differentiation, neuronal projection development, axon development, and neurotransmitter transport. Correspondingly, enriched cellular components included axons, synapses, vesicle membranes, and synaptic vesicles (**Figure 1I**). These findings support a role for TorsinA in neuronal development and function (11, 51, 52).

To compare the relative association of shared interactors with WT TorsinA and ΔE, protein abundance was normalized to the corresponding TorsinA bait abundance, thereby assigning the bait itself a WT-to-ΔE ratio of 1. Among the 137 proteins associated with both forms of TorsinA, 22 exhibited substantial differences in relative association: 17 showed reduced association and five showed increased association with ΔE compared with WT (**Figure 1J**). TorsinA-interacting protein 1 (TOR1AIP1/LAP1), an essential cofactor for TorsinA ATPase activity (18), showed markedly reduced association with ΔE. This observation is consistent with previous evidence that the ΔGAG mutation impairs interaction with its cofactors (53). Other proteins with reduced association included ATPases and ATP synthase subunits (ATP1A1, ATP2A2, and ATP5F1A), heat-shock proteins and molecular chaperones (HSP90AA1, HSPA5, and HSP90B1), the synaptic vesicle-associated protein VAT1, and several metabolic or protein-processing enzymes (PHGDH, P4HB, and GANAB). These reduced interactions may contribute to the previously reported loss-of-function properties of ΔE.

Proteins showing increased association with ΔE included factors involved in synaptic vesicle maturation and neurotransmitter release, such as UNC13B, as well as the calcium-binding chaperones calnexin (CANX) and calmegin (CLGN). These altered interactions may contribute to the functional abnormalities observed in DYT1 neurons. Most notably, exportin 1 (XPO1), a major nuclear export receptor for proteins and RNAs (54), exhibited a markedly increased association with ΔE. This finding suggested a potential molecular mechanism linking ΔE to the NCT defects previously observed in DYT1 neurons (35, 55). We therefore investigated the molecular basis of the aberrant ΔE-XPO1 interaction and its consequences for neuronal NCT and function.

### ΔE exhibits increased association with XPO1 at the nuclear envelope

We used three complementary approaches to validate the increased association between ΔE and XPO1. First, we examined the subcellular localization of endogenous TorsinA and XPO1 by immunostaining fibroblasts from patients with DYT1 dystonia and unaffected controls (**Table S1**). XPO1 was localized predominantly within the nucleus, whereas TorsinA showed a broader cytoplasmic distribution, consistent with previous reports (**Figure S2A**) (48, 56). Overlapping TorsinA and XPO1 signals were detected in both the cytoplasmic and nuclear regions (**Figure 2A**). In control cells, the overlapping signals were primarily cytoplasmic, whereas in DYT1 cells, they were enriched at the nuclear periphery. These results suggest that ΔE exhibits increased spatial proximity to XPO1 near the NE.

**Figure 2.**
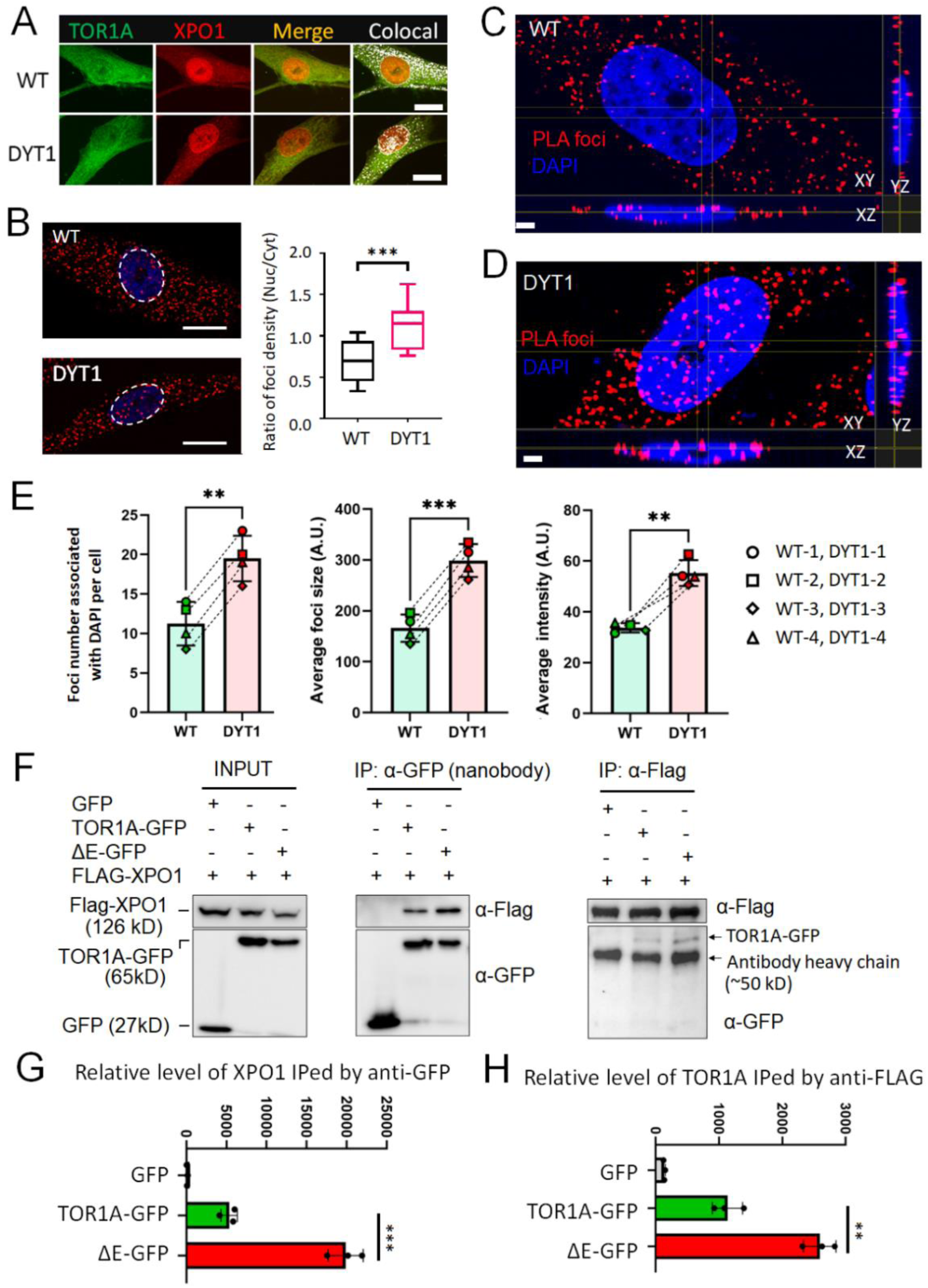
TorsinA-ΔE exhibits enhanced association with XPO1 at the nuclear envelope. **(A)** Representative confocal images of control (WT) and DYT1 patient-derived fibroblasts immunostained for endogenous TorsinA and XPO1. Colocalized signals are shown as white pixels. Scale bars, 20 µm. **(B)** Representative confocal images of proximity ligation assays (PLA) detecting the proximity of endogenous TorsinA and XPO1 in WT and DYT1 fibroblasts. The nuclear-to-cytoplasmic ratio of PLA puncta density is quantified. N (cells) = 20 for WT and 25 for DYT1 from triplicates. Scale bars, 20 µm. \*\*\**P* < 0.001; Student’s *t*-test. **(C, D)** Representative three-dimensional confocal reconstructions of PLA signals detecting endogenous TorsinA-XPO1 proximity in **(C)** WT and **(D)** DYT1 fibroblasts. Scale bars, 3 µm. **(E)** Quantification of the number, size, and fluorescence intensity of nucleus-associated PLA puncta in four pairs of DYT1 and corresponding control fibroblast lines. Twenty nuclei were analyzed per cell line. \*\**P* < 0.01 and \*\*\**P* < 0.0001; paired *t*-test. **(F)** Reciprocal Co-IP analysis of HEK cells coexpressing FLAG-tagged XPO1 with GFP, WT TorsinA-GFP, or TorsinA-ΔE-GFP. Immunoprecipitation was performed using GFP-Trap nanobody or an anti-FLAG antibody, followed by immunoblotting with the indicated antibodies. **(G, H)** Quantification of **(G)** XPO1 co-immunoprecipitated with GFP-tagged TorsinA and **(H)** GFP-tagged TorsinA co-immunoprecipitated with FLAG-XPO1. Data are from three independent experiments. \*\**P* < 0.01 and \*\*\**P* < 0.0001; Student’s *t*-test.

Second, we used a proximity ligation assay (PLA) to assess the proximity of endogenous TorsinA and XPO1 (**Figure 2B**). Few PLA puncta were detected when either the TorsinA or XPO1 primary antibody was omitted, supporting the specificity of the assay (**Figure S2B**). Compared with control cells, DYT1 fibroblasts exhibited more nucleus-associated PLA puncta and a significantly higher nuclear-to-cytoplasmic ratio of PLA signals (**Figure 2B**). High-resolution three-dimensional imaging showed that a subset of these nucleus-associated puncta localized to the nuclear periphery (**Figure 2C and D**), consistent with the NE enrichment of ΔE (**Figure 1B**). Across all four DYT1 fibroblast lines, the number, size, and fluorescence intensity of nucleus-associated PLA puncta were consistently greater than those in the corresponding controls (**Figure 2E**). These findings support an increased association between endogenous ΔE and XPO1 at or near the NE.

Third, we performed reciprocal Co-IP assays in HEK293T cells coexpressing FLAG-tagged XPO1 with GFP, WT TorsinA -GFP, or ΔE-GFP. All constructs were robustly expressed at their expected molecular weights (**Figure 2F**). Immunoprecipitation using either an anti-FLAG antibody or GFP-Trap confirmed the association between XPO1 and TorsinA and demonstrated substantially greater association of XPO1 with ΔE than with WT (**Figure 2G and H**). Together, these complementary analyses demonstrate that TorsinA and XPO1 colocalize and associate with one another and that the ΔE mutation enhances this association, particularly at or near the NE. These results independently validate the Co-IP/MS findings.

### Structural modeling predicts that the N-terminal hydrophobic segment (HS) of TorsinA engages the NES-binding cleft of XPO1

To identify regions of TorsinA that may mediate its interaction with XPO1, we used AlphaFold to generate structural models of TorsinA-XPO1 and ΔE-XPO1 complexes (**Figure 3A**) (57, 58). These models predicted that the N-terminal hydrophobic segment (HS; residues 21-50) of TorsinA contributes substantially to the interface with XPO1. Compared with WT, ΔE contained ten additional residues (within amino acids 39-50) predicted to contribute to the XPO1 interface, including five hydrophobic residues, Tyrosine 39 (Y39), Proline 40 (P40), Tyrosine 43 (Y43), Phenylalanine 46 (F46), and Alanine 47 (A47), and one positively charged residue, Arginine 41 (R41), and one negatively charged residue, Glutamic Acid 48 (E48) (**Figure 3B**).

**Figure 3.**
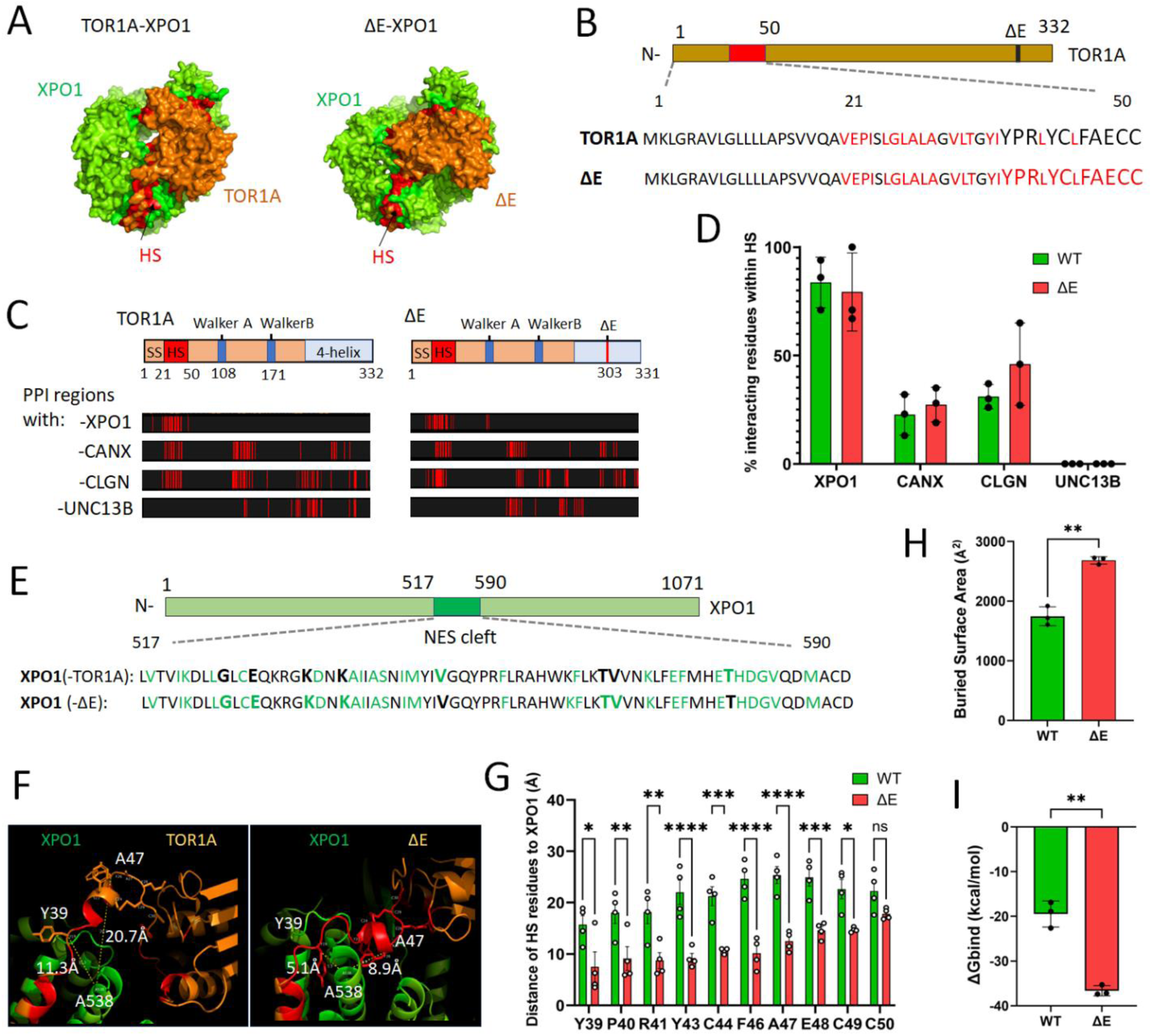
Structural modeling predicts that the N-terminal hydrophobic segment of TorsinA contributes to its interaction with XPO1. **(A)** AlphaFold-predicted structural models of WT TorsinA or TorsinA-ΔE in complex with XPO1. Predicted interface residues in TorsinA and XPO1 are shown in red and green, respectively. **(B)** Predicted contribution of the N-terminal hydrophobic segment (HS; amino acids 21–50) of TorsinA to the XPO1 interface. Compared with WT, ΔE contained additional residues (within amino acids 39-50) predicted to contribute to the interface. **(C)** Predicted TorsinA residues contributing to the modeled interfaces with XPO1, CANX, CLGN, and UNC13B. **(D)** Percentage of predicted TorsinA interface residues located within the N-terminal HS for each indicated associated protein. Values were derived from the three highest-confidence models generated for each complex. **(E)** XPO1 residues within or adjacent to the nuclear export signal (NES)-binding cleft predicted to contribute to the TorsinA interface (in green). **(F)** Enlarged views of the predicted WT TorsinA-XPO1 and TorsinA-ΔE-XPO1 interfaces. Selected HS residues in TorsinA-ΔE were positioned closer to XPO1 than the corresponding residues in WT TorsinA. **(G)** The predicated minimum distance of interface residues within HS to A538 of XPO1 based on four models with the highest confident scores. ns, not significant, * P<0.05, ** p<0.01, *** P<0.001, **** P<0.0001. Student’s T-test. **(H)** Predicted buried surface areas of the WT torsinA–XPO1 and torsinA-ΔE–XPO1 complexes, calculated from the three highest-confidence models. **(I)** Estimated binding free energies (ΔG_bind) of the WT torsinA–XPO1 and torsinA-ΔE–XPO1 complexes, calculated from the three highest-confidence models.

To examine whether the predicted contribution of the HS is specific to XPO1, we also modeled the interactions of TorsinA with CANX, CLGN, and UNC13B, three proteins that showed increased association with ΔE in the Co-IP/MS analysis (**Figure 1J**). In the TorsinA-XPO1 models, more than 65% of the TorsinA residues contributing to the predicted interface were located within the HS. By contrast, HS residues accounted for less than 36% of the predicted TorsinA contact sites in the models with CANX, CLGN, or UNC13B. No HS residues were predicted to contribute to the TorsinA-UNC13B interface, although UNC13B exhibited the greatest increase in association with TorsinA-ΔE in the proteomic analysis (**Figures 1J and 3C-D; Table S4**). These modeling results suggest that the N-terminal HS contributes preferentially to the predicted TorsinA-XPO1 interface rather than serving as a general interaction surface for all TorsinA-associated proteins.

To assess how the enhanced association of ΔE with XPO1 might affect XPO1 function, we examined the XPO1 residues contributing to the predicted interfaces. Six XPO1 residues (G526, E529, K534, K537, T563, and V564) were predicted to form contacts with ΔE but not with WT TorsinA, whereas two residues (V548 and T575) were predicted to contact WT but not ΔE (**Figure 3E**). These residues lie within or adjacent to the NES-binding cleft of XPO1, approximately spanning residues 521-605, which recognizes NES-containing cargos (59, 60). Several nearby residues, including C528, A541, K568, and F572, are established determinants of XPO1 cargo recognition, and E529 and K534 have also been reported to contact XPO1 cargos directly (60). Thus, the predicted ΔE interface overlaps the canonical cargo-binding region of XPO1, raising the possibility that aberrant ΔE binding interferes with XPO1-mediated nuclear export.

The predicted TorsinA-XPO1 interface contained numerous hydrophobic and charged residues, both of which commonly contribute to protein folding and intermolecular recognition (61–63). Examination of the modeled interfaces showed that nine out of the ten HS residues examined in ΔE were positioned closer to XPO1 than the corresponding residues in WT (**Figure 3F and G**). For example, the minimum predicted distance from Y39 and A47 to XPO1 decreased from 11.3 and 20.7 Å, respectively, in the WT complex to 5.1 and 8.9 Å in the ΔE complex (**Figure 3F**). Moreover, the predicted buried surface area of the ΔE-XPO1 complex was greater than that of the WT-XPO1 complex (**Figure 3H**), consistent with a more extensive protein interface (64). Computational estimation of binding free energy also yielded a more favorable ΔG_bind for the ΔE complex than for the WT complex (65) (**Figure 3I**). Collectively, these structural analyses predict that the ΔE mutation promotes a more extensive and energetically favorable association with XPO1 involving the N-terminal HS.

To determine whether the predicted enhancement of XPO1 association is specific to the ΔE mutation, we modeled XPO1 interactions with additional TorsinA variants. These included K108A and E171Q, which disrupt ATP binding and hydrolysis, respectively (18, 45), as well as disease-associated variants identified in individuals with dystonia: E121K, D194V, and R288Q, associated with early-onset dystonia (66, 67), and F205I, associated with late-onset focal dystonia (68) (**Figure S3A**). In all modeled complexes, most TorsinA residues contributing to the predicted XPO1 interface were located within the N-terminal HS (**Figure S3B** and **C; Table S5**). However, among the variants examined, only ΔE displayed both an increased predicted buried surface area and a more favorable estimated ΔG_bind relative to WT TorsinA (**Figure S3D** and **E**). This result is consistent with previous observations that ΔE, but not the other examined variants, produces prominent redistribution of TorsinA to the NE (67). Together, these computational analyses predict that the N-terminal HS contributes to TorsinA-XPO1 association and that the ΔE mutation uniquely strengthens this interaction.

### Deletion of the N-terminal HS reduces ΔE association with XPO1 and alters its subcellular distribution

To experimentally test the AlphaFold-based prediction, we generated constructs encoding WT TorsinA or ΔE lacking the N-terminal HS (ΔHS) and examined their association with XPO1 (**Figure 4A**). HEK293T cells were cotransfected with constructs expressing FLAG-tagged XPO1 and GFP-tagged WT TorsinA, ΔE, or their corresponding ΔHS variants. All proteins were expressed at their expected molecular weights (**Figure 4B**, input). Co-IP analysis showed that HS deletion significantly reduced the association of both WT and ΔE with XPO1 (**Figure 4B** and **C**), supporting a role for the N-terminal HS in the TorsinA-XPO1 interaction. Notably, HS deletion produced a greater reduction in the amount of XPO1 co-immunoprecipitated with ΔE than with WT (approximately 75% versus 50%, respectively). These findings indicate that although the HS contributes to the association of both forms of TorsinA with XPO1, ΔE shows a greater dependence on this region. The reduction observed with WT TorsinA further raises the possibility that the TorsinA-XPO1 interaction has a physiological role in regulating XPO1-dependent nuclear transport.

**Figure 4.**
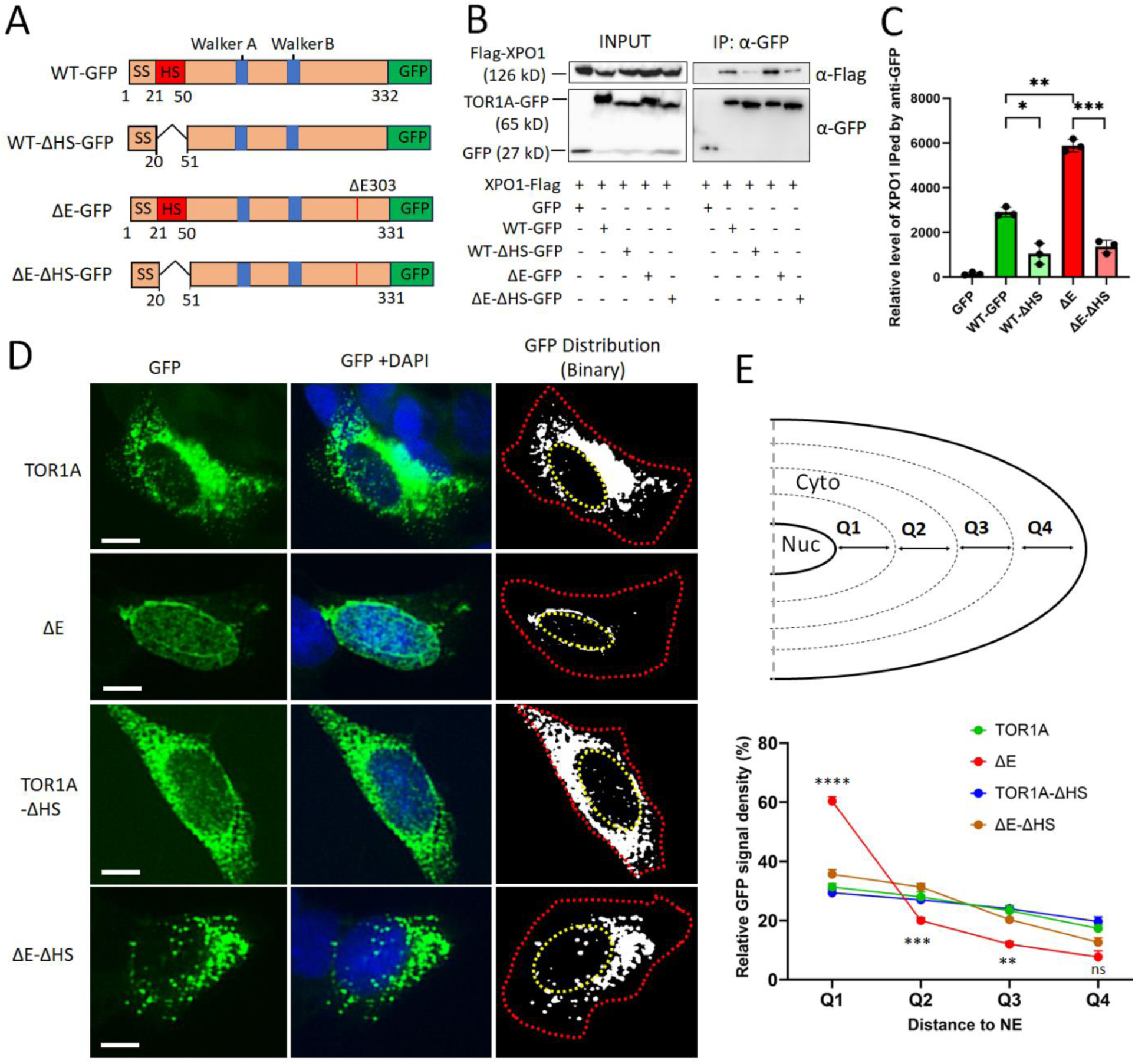
Deletion of the N-terminal hydrophobic segment reduces TorsinA-ΔE association with XPO1 and alters its subcellular distribution. **(A)** Schematic of GFP-tagged WT TorsinA and TorsinA-ΔE constructs with or without deletion of the N-terminal HS (ΔHS). SS, signal sequence; HS, hydrophobic segment. **(B)** Co-immunoprecipitation analysis of GFP-tagged TorsinA variants in HEK cells coexpressing FLAG-tagged XPO1. GFP-tagged proteins were immunoprecipitated using GFP-Trap and analyzed by immunoblotting with the indicated antibodies. **(C)** Quantification of co-immunoprecipitated FLAG-XPO1 normalized to the corresponding GFP-tagged TorsinA bait. Data are from three independent experiments. \**P* < 0.05, \*\**P* < 0.01, and \*\*\**P* < 0.001; one-way ANOVA followed by Dunnett’s multiple-comparisons test. **(D)** Representative fluorescence images of SH-SY5Y cells expressing GFP-tagged WT TorsinA, ΔE, or their corresponding ΔHS variants. GFP fluorescence is also displayed in grayscale to facilitate visualization of its subcellular distribution. Red dotted lines delineate the cell boundaries, and yellow dotted lines delineate the nuclei. Scale bars, 10 µm. **(E)** Schematic of the image-analysis strategy and quantification of the cytoplasmic distribution of the GFP-tagged TorsinA variants shown in **(D)**. N (cells) = 45 of each condition from three independent experiments. Compared ΔE-ΔHS to ΔE, ns, no significant difference; **p<0.01; ***p<0.0001. Student’s *T*-test.

We next investigated whether HS deletion affected the subcellular distribution of TorsinA. SH-SY5Y neuroblastoma cells were transduced with lentiviral vectors expressing GFP-tagged WT TorsinA, ΔE, or their respective ΔHS variants. SH-SY5Y cells were selected because their relatively large cell bodies facilitated visualization of subcellular protein distribution. Consistent with our observations in induced human neurons (**Figure 1B**), WT TorsinA exhibited a broad ER-associated cytoplasmic distribution, whereas ΔE was prominently enriched at the NE (**Figure 4D**). Deletion of the HS markedly redistributed ΔE from the NE to a broader cytoplasmic pattern but produced no apparent change in the distribution of WT TorsinA (**Figure 4D and E**). Together, these results show that the N-terminal HS contributes to both the enhanced association of ΔE with XPO1 and its enrichment at the NE.

### Deletion of the N-terminal HS restores nuclear export in ΔE expressing cells

Because the HS contributes to the enhanced association of ΔE with XPO1 at the NE, we hypothesized that deleting the HS would alleviate XPO1 dysfunction and restore nuclear export. To test this hypothesis, we assessed nuclear export in SH-SY5Y cells co-transduced with lentiviral vectors expressing GFP-tagged TorsinA variants and an RFP-based nuclear export reporter containing a nuclear export signal (RFP-NES), which has been validated XPO1-dependent (36, 38, 69). Impaired nuclear export increases the nuclear retention of RFP-NES and therefore produces a higher nuclear-to-cytoplasmic (N/C) fluorescence ratio (**Figure 5A**).

**Figure 5.**
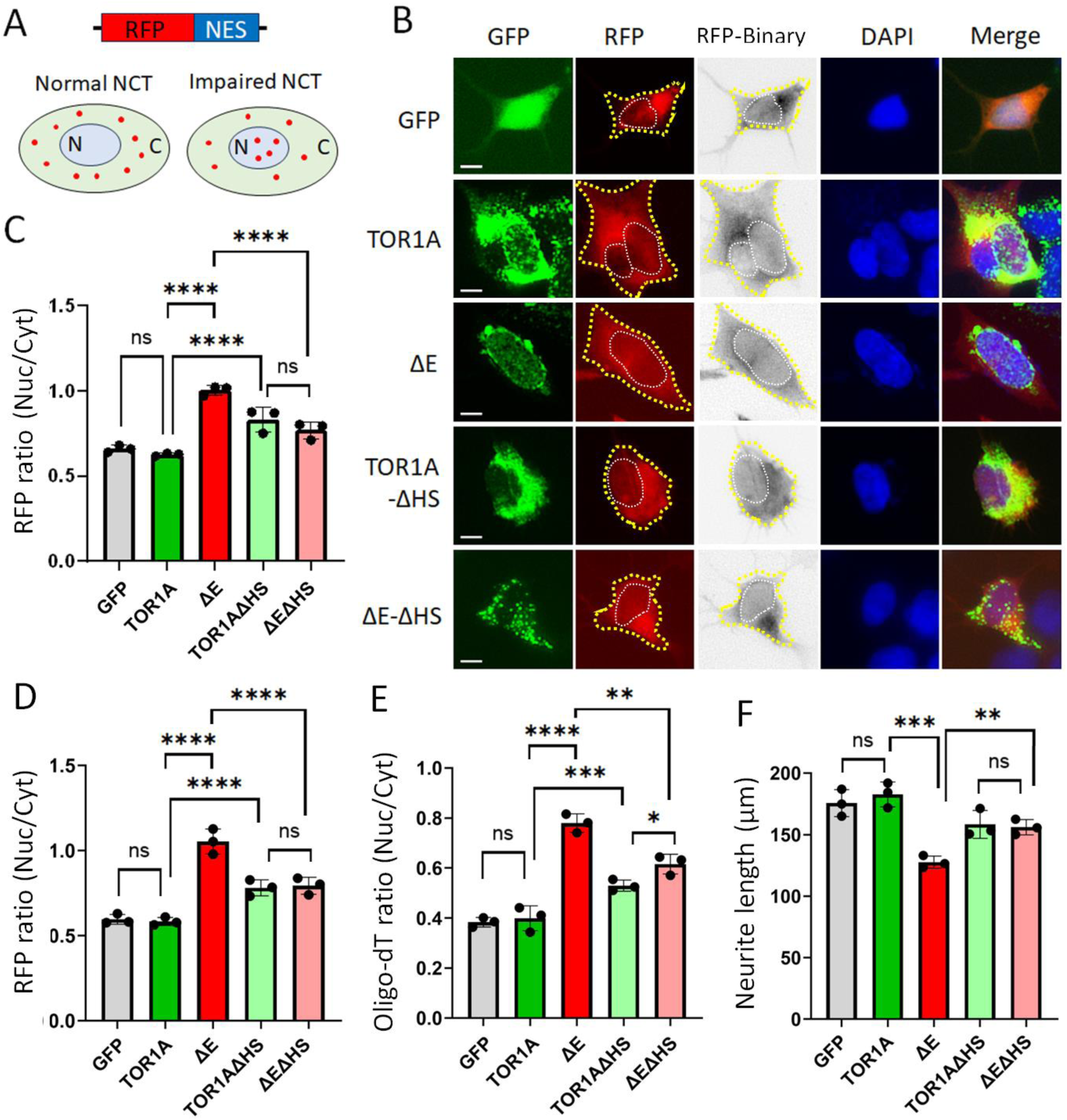
Deletion of the N-terminal hydrophobic segment alleviates nuclear export defects in ΔE-expressing cells. **(A)** Schematic of the nuclear export reporter comprising red fluorescent protein fused to a nuclear export signal (RFP-NES). **(B)** Representative fluorescence images of SH-SY5Y cells expressing RFP-NES together with GFP, WT TorsinA, ΔE, or their corresponding ΔHS variants at 3 dpi. Nuclei and cell boundaries are outlined by white and yellow dashed lines, respectively. RFP-NES distribution is displayed as both red fluorescence and thresholded binary images. Scale bar, 10 µm. **(C)** Quantification of the nuclear-to-cytoplasmic (N/C) ratio of RFP-NES fluorescence intensity in the undifferentiated SH-SY5Y cells shown in **(B)**. **(D)** Quantification of the RFP-NES N/C fluorescence ratio in differentiated SH-SY5Y cells expressing GFP or the indicated TorsinA variants at 7 dpi. **(E)** Quantification of the N/C ratio of poly(A)+ RNA fluorescence detected by oligo(dT) fluorescence in situ hybridization (FISH) in differentiated SH-SY5Y cells at 7 dpi. **(F)** Quantification of relative neurite length in differentiated SH-SY5Y cells expressing GFP or the indicated TorsinA variants at 5 dpi. For **(C–F)**, each point represents the mean of more than 50 cells from one independent experiment. N = 3 independent experiments. Data are presented as mean ± SD. ns, not significant; \**P* < 0.05, \*\**P* < 0.01, \*\*\**P* < 0.001, and \*\*\*\**P* < 0.0001; one-way ANOVA followed by Dunnett’s multiple-comparisons test.

In cells expressing GFP or WT TorsinA, RFP-NES was localized predominantly in the cytoplasm, resulting in a low N/C ratio. By contrast, cells expressing ΔE exhibited significantly greater nuclear accumulation of RFP-NES and a correspondingly higher N/C ratio, indicating impaired nuclear export and confirming our previous findings (35, 36) (**Figure 5B and C**). Deletion of the HS from ΔE significantly reduced the RFP-NES N/C ratio relative to that in cells expressing ΔE, demonstrating restoration of nuclear export activity (**Figure 5B and C**). Interestingly, cells expressing WT TorsinA-ΔHS exhibited a significantly higher RFP-NES N/C ratio than cells expressing full-length WT TorsinA, suggesting that HS deletion impairs nuclear export in the WT context. Because HS deletion also reduced the association between WT TorsinA and XPO1 (**Figure 4B and C**), these findings raise the possibility that an appropriately regulated TorsinA-XPO1 association contributes to XPO1-mediated nuclear export under physiological conditions. In contrast, the abnormally enhanced association of ΔE with XPO1 may interfere with this process.

We next examined nuclear export in SH-SY5Y-derived neurons differentiated under defined culture conditions (35). Consistent with the results in undifferentiated cells, deletion of the HS from ΔE reduced the nuclear retention of both the RFP-NES protein reporter and poly(A)+ RNA, as assessed by fluorescence in situ hybridization (**Figure 5D and E**). HS deletion also improved neurite outgrowth in ΔE-expressing cells (**Figure 5F**), linking restoration of NCT to an improvement in neuronal morphology. The rescue of poly(A)+ RNA export by HS deletion was less pronounced than that of RFP-NES export, consistent with the direct role of XPO1 in exporting NES-containing proteins and the involvement of distinct transport pathways in bulk mRNA export (70, 71). Together, these findings demonstrate that deletion of the N-terminal HS alleviates protein-and RNA-export defects in ΔE-expressing cells. The results support a model in which the aberrant HS-dependent association of ΔE with XPO1 compromises nuclear export.

### Deletion of the N-terminal HS from ΔE improves neurite outgrowth and maturation-associated gene expression in induced human neurons

To determine whether deletion of the HS from ΔE ameliorates neuronal deficits, we transduced hiPSC-derived MNs with lentiviral vectors expressing GFP-tagged WT TorsinA, ΔE, or their corresponding ΔHS variants. Consistent with previous reports (35, 36), neurons expressing ΔE exhibited shorter neurites and fewer primary and secondary branches than neurons expressing WT TorsinA at 7 dpi, an early stage of neuronal development (**Figure 6A-D**). Expression of ΔE-ΔHS significantly increased neurite length and branching relative to ΔE, with a greater improvement in secondary than in primary branches (**Figure 6A-D**).

**Figure 6.**
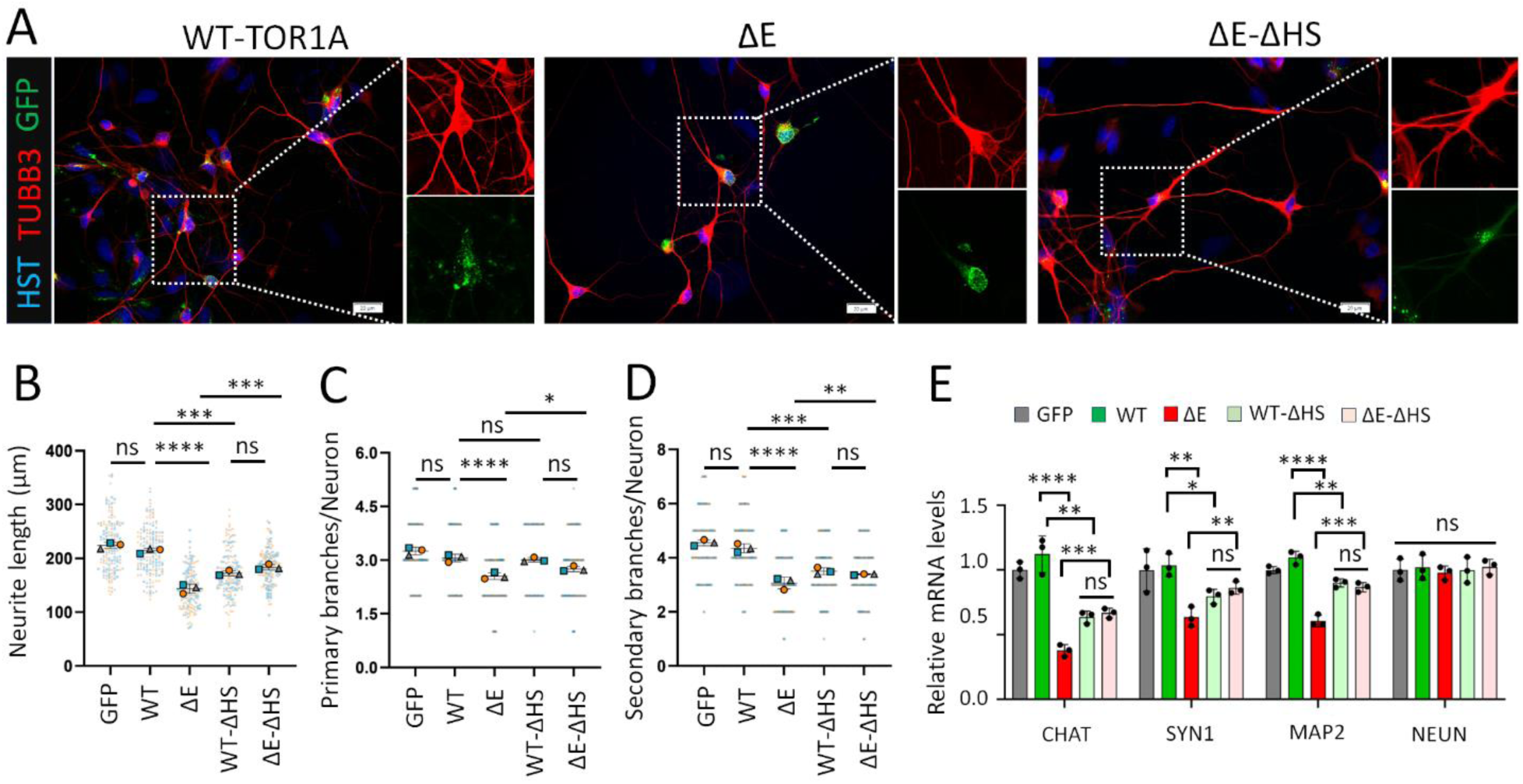
Deletion of the N-terminal hydrophobic segment from TorsinA-ΔE improves neurite outgrowth and maturation-associated gene expression in induced human neurons. **(A)** Representative fluorescence images of hiPSC-derived motor neurons expressing GFP-tagged WT TorsinA, ΔE, or their corresponding ΔHS variants at 7 dpi. Scale bars, 20 µm. **(B–D)** Quantification of **(B)** relative neurite length, **(C)** number of primary branches, and **(D)** number of secondary branches in hiPSC-derived motor neurons expressing the indicated TorsinA variants at 7 dpi. A total of 150 neurons per group were analyzed across three independent experiments, with data from each experiment shown in a distinct color. **(E)** RT-qPCR analysis of the indicated neuronal maturation-associated genes in hiPSC-derived motor neurons expressing the TorsinA variants at 14 dpi. Data are from three independent experiments. Data are presented as mean ± SD ns, not significant; \**P* < 0.05, \*\**P* < 0.01, \*\*\**P* < 0.001, and \*\*\*\**P* < 0.0001; one-way ANOVA followed by Dunnett’s multiple-comparisons test.

Gene-expression analysis at 14 dpi further showed that neurons expressing ΔE-ΔHS had significantly higher transcript levels of several neuronal maturation-associated genes, including *CHAT*, *SYN1*, and *MAP2*, than neurons expressing ΔE (**Figure 6E**). By contrast, *RBFOX3* (NeuN) expression was not significantly altered. Because these genes participate in processes associated with neuronal maturation, including neurotransmitter synthesis, synapse formation, and dendritic development, these results indicate that HS deletion alleviates the neurite outgrowth defects caused by ΔE and partially restores maturation-associated gene expression.

### Depletion of the TorsinA cofactors LAP1 or LULL1 does not alter TorsinA-XPO1 association

TorsinA forms complexes with cofactors LAP1 and LULL1 (18), which are required for its ATPase activity. To determine whether these cofactors contribute to the association between TorsinA and XPO1, we used two complementary approaches. First, we examined whether LAP1 or LULL1 was present in anti-FLAG immunoprecipitates from cells expressing FLAG-tagged XPO1. Neither cofactor was detected in the immunoprecipitated fractions (data not shown), providing no evidence that LAP1 or LULL1 is part of the TorsinA-XPO1 complex under these experimental conditions. Second, we generated lentiviral vectors expressing shRNAs targeting LAP1 or LULL1 and confirmed efficient depletion of the corresponding proteins (**Figure S4A**). Co-IP assays consistently showed greater XPO1 association with ΔE than with WT TorsinA. Depletion of either LAP1 or LULL1 did not significantly alter the association of WT TorsinA or ΔE with XPO1 (**Figure S4**). Together, these results indicate that LAP1 and LULL1 are individually dispensable for the TorsinA-XPO1 association under the conditions examined.

### Structural modeling predicts enhanced XPO1 association with ΔE-containing TosinA dimers

Because the disease-associated ΔGAG mutation is typically present in the heterozygous state, both WT TorsinA and ΔE are expressed in cells from individuals with DYT1 dystonia (6). We therefore used AlphaFold to model XPO1 in complex with WT-WT and ΔE-ΔE homodimers or a WT-ΔE heterodimer. The models predicted a greater number of interface residues between XPO1 and ΔE-containing TorsinA dimers than between XPO1 and the WT-WT homodimer (**Figure S5A-C**). Moreover, the predicted buried surface area progressively increased from the WT-WT homodimer to the WT-ΔE heterodimer and then to the ΔE-ΔE homodimer (**Figure S5D**). This pattern was accompanied by decreasingly estimated binding free energies (ΔG_bind) (**Figure S5E**). These computational results predict that the presence of ΔE strengthens the association of TorsinA dimers with XPO1, including in the disease-relevant WT-ΔE heterodimeric configuration.

### Expression of HS-derived peptides ameliorates cellular deficits in DYT1 neurons

Because the N-terminal HS contributes to the enhanced association of ΔE with XPO1, and HS deletion alleviates nuclear export defects, we hypothesized that HS-derived peptides could interfere with the aberrant ΔE-XPO1 interaction. To test this hypothesis, we generated lentiviral constructs expressing GFP-tagged HS peptides of different lengths, selected to compare their activity and solubility. These included a peptide containing residues predicted to preferentially contribute to the ΔE-XPO1 interface (amino acids 39-50), as well as peptides containing residues predicted to contribute to the interfaces of both WT and ΔE with XPO1 (amino acids 21-50 and 21-35) (**Figure 7A**). The GFP-tagged peptides were expressed at their expected molecular weights and at levels comparable to those of the GFP control (**Figure 7B**).

**Figure 7.**
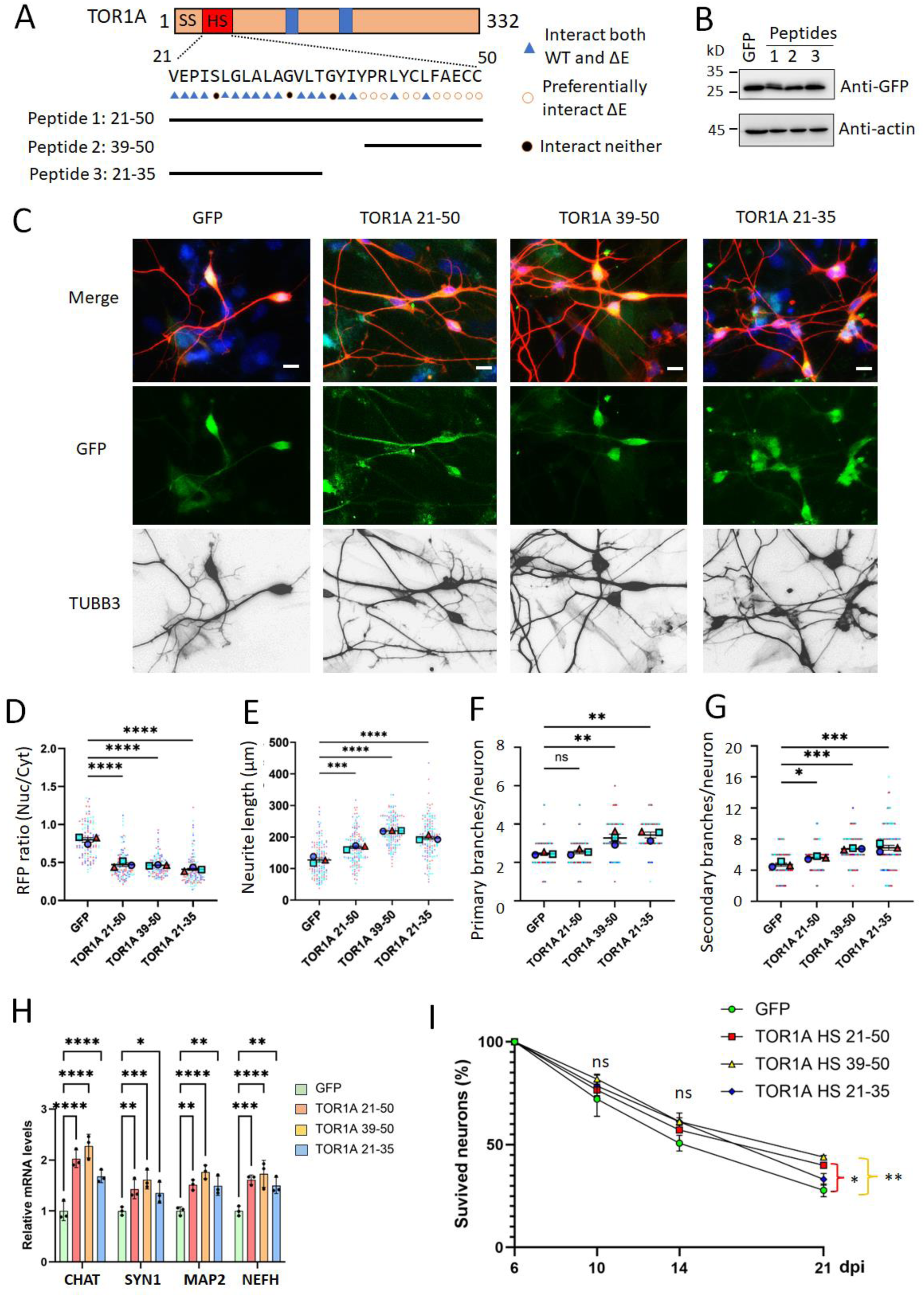
Expression of TorsinA HS-derived peptides ameliorates cellular deficits in DYT1 motor neurons. **(A)** Schematic of the GFP-tagged peptides derived from the N-terminal hydrophobic segment (HS) of TorsinA. **(B)** Immunoblot analysis of DYT1 hiPSC-derived motor neurons transduced with lentiviral vectors expressing GFP or the indicated GFP-tagged HS-derived peptides. **(C)** Representative fluorescence images of DYT1 hiPSC-derived motor neurons expressing GFP or the indicated GFP-tagged HS-derived peptides. Motor neurons were cocultured with astrocytes and imaged at 6 dpi. Scale bars, 20 µm. **(D)** Quantification of nuclear export using the RFP-NES reporter in DYT1 hiPSC-derived motor neurons at 6 dpi. Nuclear export activity was assessed by measuring the nuclear-to-cytoplasmic ratio of RFP-NES fluorescence. **(E)** Quantification of neurite length in DYT1 hiPSC-derived motor neurons at 6 dpi. **(F, G)** Quantification of the numbers of **(F)** primary and **(G)** secondary branches in DYT1 hiPSC-derived motor neurons at 10 dpi. For **(D-G)**, 100 neurons per group were analyzed across three independent experiments, with data from each experiment shown in a distinct color. ns, not significant; \**P* < 0.05, \*\**P* < 0.01, and \*\*\**P* < 0.001; one-way ANOVA followed by Dunnett’s multiple-comparisons test. **(H)** RT-qPCR analysis of the indicated neuronal maturation-associated genes in DYT1 hiPSC-derived motor neurons at 14 dpi. Data are from three independent experiments. \**P* < 0.05, \*\**P* < 0.01, \*\*\**P* < 0.001, and \*\*\*\**P* < 0.0001; one-way ANOVA followed by Dunnett’s multiple-comparisons test. **(I)** Survival of DYT1 hiPSC-derived motor neurons expressing GFP or the indicated GFP-tagged HS-derived peptides. The number of surviving neurons at 6 dpi was defined as 100% for each condition, and subsequent measurements were normalized to this baseline. A total of 200 neurons per group were analyzed across three independent experiments. ns, not significant; \**P* < 0.05 and \*\**P* < 0.01 versus the GFP control; one-way ANOVA followed by Dunnett’s multiple- comparisons test.

We systematically evaluated the effects of these HS-derived peptides in MNs differentiated from DYT1 patient-derived iPSC lines and their corresponding isogenic controls (**Figures 7C and S6**) (72, 73). Compared with GFP alone, expression of the HS-derived peptides significantly improved nuclear export in DYT1 MNs (**Figure 7D**). Peptide expression also promoted neurite outgrowth (**Figure 7E**), increased the numbers of primary and secondary branches (**Figure 7F and G**), and elevated the expression of neuronal maturation-associated genes (**Figure 7H**). At 21 dpi, expression of peptides 1 (amino acids 21-50) and 2 (amino acids 39-50) also significantly increased DYT1 MN survival (**Figure 7I**). By contrast, the peptides produced only modest effects in isogenic control neurons (**Figure S6**). Among the peptides examined, peptide 2, comprising amino acids 39-50, produced the strongest overall improvement across the measured outcomes. This result is consistent with the structural models predicting that this region contributes preferentially to the ΔE-XPO1 interface. Together, these findings demonstrate that expression of HS-derived peptides, particularly the amino acid 39-50 peptide, ameliorates nuclear export, morphological, maturation-associated, and survival deficits in DYT1 MNs. The results further identify the HS-mediated ΔE-XPO1 interaction as a potentially targetable disease mechanism.

## Discussion

In this study, we identify an aberrant interaction between TorsinA-ΔE and XPO1 as a mechanism contributing to nucleocytoplasmic transport dysfunction in DYT1 dystonia. Proteomic analysis revealed increased association of ΔE with XPO1, which was supported by colocalization, proximity ligation, and reciprocal co-immunoprecipitation assays. Structural modeling predicted that the N-terminal hydrophobic segment of TorsinA engages a region within or adjacent to the NES-binding cleft of XPO1 and that the ΔE mutation strengthens this interface. Deletion of the HS reduced ΔE association with XPO1, altered its NE enrichment, and alleviated nuclear export and neuronal developmental deficits. Importantly, HS-derived peptides improved nuclear export, morphology, maturation-associated gene expression, and survival in patient-derived DYT1 motor neurons. Together, these findings support an HS-dependent gain-of-function mechanism that complements the established loss-of-function effects of the ΔE mutation and identifies the ΔE-XPO1 interaction as a potential therapeutic target.

Although ΔE has traditionally been viewed as a loss-of-function variant (8), our findings suggest that its pathogenic activity is more complex. The reduced association of ΔE with LAP1 and multiple other proteins is consistent with loss of normal TorsinA functions. In parallel, however, ΔE exhibited a markedly enhanced association with XPO1. Thus, the ΔE mutation may produce simultaneous loss of physiological interactions and gain of an aberrant interaction. Its accumulation at the NE could increase local access to XPO1 and promote aberrant engagement of the nuclear export machinery. Loss and gain of function are not mutually exclusive (74, 75). This dual mechanism may help explain why the ΔE mutation produces cellular effects that cannot be attributed solely to reduced TorsinA activity.

TorsinA has been shown to play an important role in maintaining NE structure and function. Loss or dysfunction of torsin proteins causes nuclear membrane abnormalities, including NE deformation, membrane blebbing, and impaired nuclear pore complex assembly or maturation (3, 9, 34, 76). These findings have established a close relationship among torsin activity, NE homeostasis, and nuclear pore biogenesis. However, how the DYT1-associated ΔE mutation translates these structural abnormalities into impaired nuclear transport and neuronal dysfunction has remained incompletely understood. Our previous studies demonstrated that DYT1 patient-derived neurons exhibit defective NCT, including nuclear retention of poly(A)+ RNA and altered localization of protein-transport reporters (35, 36, 43). These defects were accompanied by NE abnormalities, nuclear Lamin B1 mislocalization, impaired neurite outgrowth, and delayed neuronal maturation, suggesting that disruption of NE organization and NCT represents an important component of DYT1 pathogenesis. One possible explanation is that defective TorsinA activity impairs nuclear pore biogenesis or maturation, thereby broadly reducing transport across the NE (34, 35). The present study extends this model by identifying XPO1 as a transport receptor that associates aberrantly with ΔE. Thus, TorsinA dysfunction may compromise NCT through at least two potentially complementary mechanisms: disruption of NE/NPC organization and inappropriate engagement of the nuclear transport machinery.

XPO1 is a major nuclear export receptor that recognizes leucine-rich NES and transports a diverse group of protein cargos, including transcription factors, signaling molecules, cell-cycle regulators, and proteins involved in stress responses and protein homeostasis (59, 77, 78). Consequently, abnormal XPO1 localization or availability could alter multiple cellular pathways simultaneously, even without causing complete inhibition of nuclear export. Our results show that ΔE exhibits increased association and spatial proximity with XPO1, particularly at or near the NE, and that expression of ΔE increases the nuclear retention of an NES-containing reporter. These observations support a model in which aberrant engagement of XPO1 by ΔE reduces the efficiency of XPO1-dependent cargo export. The predicted localization of the ΔE interface within or adjacent to the NES-binding cleft of XPO1 further raises the possibility that ΔE interferes with cargo recognition or export-complex formation, although direct competition with NES-containing cargos remains to be experimentally established.

The effects of ΔE may also extend beyond XPO1-dependent protein export. Bulk poly(A)+ RNA export is mediated predominantly by the NXF1-NXT1 pathway rather than directly by XPO1 (38, 79). Therefore, the poly(A)+ RNA-export defect observed in DYT1 neurons likely reflects broader disruption of the NE, NPCs, or interconnected transport pathways. Consistent with this interpretation, deletion of the TorsinA N-terminal HS produced a stronger rescue of NES-reporter export than of poly(A)+ RNA export. Together, these findings position the ΔE-XPO1 interaction within a broader model in which the ΔE mutation perturbs nuclear transport both through abnormal engagement of a specific export receptor and through more general disruption of NE and nuclear pore homeostasis. Because developing neurons depend on tightly regulated nucleocytoplasmic trafficking to coordinate transcriptional programs, stress responses, neurite growth, and synaptic maturation, these combined transport defects may contribute directly to the developmental neuronal phenotypes associated with DYT1 dystonia.

The ability of HS-derived peptides to ameliorate multiple disease-associated phenotypes has important therapeutic implications. Expression of these peptides improved nuclear export, neurite outgrowth and maturation in DYT1 neurons while producing only modest effects in isogenic control neurons. These findings suggest that the aberrant ΔE-XPO1 interaction may be selectively disrupted without broadly suppressing the physiological functions of XPO1. This distinction is important because global inhibition of XPO1 would be expected to affect numerous essential cargos and could cause substantial toxicity. Targeting the mutation-enhanced protein-protein interaction may therefore offer greater specificity than directly inhibiting XPO1-mediated transport. Among the sequences examined, the peptide encompassing TorsinA residues 39-50 produced the most consistent rescue across the measured endpoints. This region contains several residues predicted to contribute preferentially to the ΔE-XPO1 interface, supporting its further development as a lead therapeutic sequence. The present peptides were expressed intracellularly as GFP-tagged constructs and therefore serve primarily as proof-of-concept tools. Future development could focus on shorter and more stable peptides, cell-penetrating peptides, peptidomimetics, or small molecules that reproduce the relevant structural and chemical properties of this region. Optimization should seek to preserve disruption of the pathological ΔE-XPO1 association while minimizing interference with the physiological association between WT TorsinA and XPO1. The limited effects observed in isogenic control neurons are encouraging, but detailed dose-response, target-engagement, toxicity, and off-target analyses will be required to define a therapeutic window.

An additional mechanistic question concerns the apparent interaction between TorsinA and XPO1 across their conventionally assigned subcellular compartments. Canonical TorsinA is localized within the ER/NE lumen (18), whereas XPO1 functions primarily on the nucleoplasmic and cytoplasmic faces of the nuclear pore (59). The physical basis for their association therefore remains unresolved. Potential explanations include the existence of a noncanonical cytosolic pool of TorsinA (22, 23), stress-dependent redistribution (20, 21), altered membrane insertion or topology of ΔE, or association through intermediary proteins. This explanation is supported by recent studies that Torsin ATPase activity and the coactivator are largely dispensable for the fly fat body lipidome and germ cell development (24, 25). Defining the membrane topology and subcellular pool responsible for the interaction will be essential for establishing the mechanism and determining where a therapeutic inhibitor must act.

## Materials and Methods

### Cell lines and culture condition

The use of human cell lines, including human induced pluripotent stem cells (hiPSCs), in this study was approved by the Institutional Review Board (IRB) of Louisiana State University Health Sciences Center at Shreveport (LSUHSC-S, IRB ID: STUDY00002945). All cell lines used in this study were listed in Supplemental **Table S1**. HEK 293T cells (CRL-11268) and SH-SY5Y (CRL-11266) cells were purchased from ATCC. DYT1 fibroblast cell lines and sex- and age-matched healthy controls were obtained from the *Coriell Institute for Medical Research* or *NINDS Human Cell and Data Repository* (NHCDR). Human induced pluripotent stem cell (hiPSC) lines have been generated in our previous studies (35, 72, 73). All hiPSCs were maintained in complete mTeSR1 medium (STEMCELL Technologies) on Matrigel (Corning) coated dishes at 37 °C and 5% CO_2_ with saturating humanity, and the medium was daily replaced. The recipes for different media were described as supplemental information.

### Plasmid construction and lentivirus production

A third-generation lentiviral vector (*pCSC-SP-PW-IRES-GFP*) was used to express reprogramming factors NGN2-IRES-ISL1-T2A-LHX3 (Addgene #221814) as previously reports (36, 41). The same lentiviral vector was used to express GFP-tagged TorsinA, TorsinA variants, and hydrophobic peptides. Another third-generation lentiviral vector (*LV-CAG-mCherry-miRE-Luc*) was used to express shRNAs targeting TOR1AIP1 (LAP1) and TOR1AIP2 (LULL1) as described in previously reports (35, 36, 80). The targeting sequences (gRNAs) were listed in Supplemental **Table S2**. The dual reporter 2Gi2R (Addgene #71396) was generously provided by Dr. Fred H. Gage (81). The transport reporters of RFP-NES (Addgene #221812) and RFP-NLS (Addgene #221813) were constructed as previously reported (36). All plasmids were verified by restriction enzyme digestions and DNA sequencing. Each of these vectors was co-transfected with packaging plasmids (Addgene #12251, #12253, and #12259) into HEK293T cells for virus production (82, 83). Replication-incompetent lentiviruses were produced, and viral supernatants were collected at 48 hrs and 72 hrs post-transfection as previously described (37, 84). The viral supernatants were filtered through 0.45 μm syringe filters and stored at 4 °C prior to cell transduction.

### Generation of hiPSC-derived MNs

MNs were prepared from hiPSCs as previously described (39–41). Briefly, hiPSCs were cultured in mTeSR1 medium with 10 µM all-trans-retinoic acid (RA, Sigma) and 0.5 mM VPA in Matrigel-coated 6-well plates for 7 days. Cells were then digested with Versene and gently pipetted into small clumps supplemented with 10 μM Y-27632 (STEMCELL Technologies). Cell clumps were cultured in KOSR medium for 4 days, followed by cultured in NSP medium for another week. The neurospheres were then dissociated into single cells with Accutase (Innovative Cell Technologies) and maintained in neural progenitor cell medium. For MN differentiation (41, 42, 72), neural progenitor cells were plated into Matrigel-coated plates at a density of 3×10^4^ cells/cm^2^ and transduced with a lentiviruse expressing *NEUROG2-IRES-ISL1-T2A-LHX3*. Culture medium was replaced the next day with neuronal maturation medium. Neurons were dissociated with Accutase on day 5 and replated onto Matrigel-coated coverslips with or without the presentence of astrocytes depending on desired experiments. The medium was half changed twice a week until analysis.

### Generation of SH-SY5Y-derived neurons

The SH-SY5Y neuroblastoma cells were cultured on gelatin-coated plates and maintained in SH-SY5Y medium. To induce neuronal differentiation, the media was switched to SH-SY5Y differentiation medium. The medium was half changed every other day. SY5Y cells gradually became neuron-like morphology within one week. To ectopically overexpress genes in SY5Y-derived neurons, transduced lentiviruses expressing genes of interest when cells reached about 50% confluence. After an overnight exposure to lentivirus, the media was replaced the next day with SH-SY5Y differentiation medium, and half changed twice a week until analysis.

### Co-immunoprecipitation coupled with mass spectrometry (Co-IP/MS) assay

Cultured neurons were harvested and lysed using a lysis buffer containing 50 mM Tris-HCl (pH 7.4), 150 mM NaCl, 0.5 mM EDTANa2, 2% Triton-X 100, 1mM PMSF, and DNase I and RNase A. Anti-GFP Nanobody coupled with agarose beads (Proteintech, gta) was used to precipitate GFP and GFP tagged proteins as previously described (43, 85–87). For LC-MS/MS analysis, ∼10 µg of protein from each sample was loaded into 12% SDS-PAGE, and then stained with Imperial Protein Stain (ThermoFisher). The Gel lanes of each sample were cut into 2mm X 2mm cubes, and then performed trypsin digestion. The digested peptides were vacuum-dried and resolved in the LC/MS grade water with 0.1% (v/v) formic acid, followed by untargeted discovery proteomics analysis.

This LC-MS/MS analysis was carried out using an Ultimate 3000 RSLCnano system connected to an Orbitrap Exploris 480 mass spectrometer. The digested peptides (5.0 µL) were loaded onto a trap column (PepMap C18, 2 cm × 100 μm, 100 Å) at a flow rate of 20 μL/min using 0.1% formic acid, and separated on an analytical column (EasySpray 50 cm × 75 μm, C18 1.9 μm, 100 Å) with a flow rate of 300 nL/min with a linear gradient of 5 to 45% solvent B (100% ACN, 0.1% formic acid) over a 120 min gradient. Both precursor and fragment ions were acquired in the Orbitrap mass analyzer. Precursor ions were acquired in m/z range of 375-1500 with a resolution of 120,000 (at m/z 200). Precursor fragmentation was carried out using the higher-energy collisional dissociation method using normalized collision energy (NCE) of 32. The fragment ions were acquired at a resolution of 150,000 (at m/z 200). The scans were arranged in top-speed method with 3 sec cycle time between MS and MS/MS. Ion transfer capillary voltage were maintained at 2.1 kV.

The raw Mass Spec data were analyzed using Proteome Discover (version 2.5, Thermo Fisher Scientific) software package with SequestHT using species-specific fasta database and the Percolator peptide validator. Cysteine alkylation was set as a fixed modification, and deamidation of asparagine residues was selected as variable modifications. The abundance of GFP peptides in each group (GFP, TorsinA-GFP, and ΔE-GFP) served as an internal loading control to normalize the abundance of identified proteins. Specific binding was defined by a threshold of at least 5-fold enrichment in TorsinA-GFP, or ΔE-GFP compared to GFP controls. To acquire protein-protein interaction networks and their functional clusters, network analysis was performed using an online database (https://string-db.org) and Gene Ontology (GO) analysis was conducted using PANTHER (protein annotation through evolutionary relationship) classification system (http://www.pantherdb.org/) (88).

### Western blotting analysis

Cells were lysed in lysis buffer composed of 50 mM Tris-HCl buffer (pH 8.0), 150 mM NaCl, 1% NP40, 1% Triton X-100, 0.1% SDS, 0.5% sodium deoxycholate, and protease inhibitor cocktail (Roche). Equal amounts of cell lysates (20 µg per lane) were used for SDS-PAGE and western blot analysis as previously described (35, 87, 89). Primary antibodies and dilutions were listed in Supplemental **Table S3.** HRP-conjugated secondary antibodies and Clarity Western ECL substrate (Bio-Rad) were used to visualize the protein bands with a BioRad ChemiDoc.

### Protein structure and interface analysis

Protein structures and protein-protein interactions were predicted using AlphaFold (58, 90). Briefly, the amino acid sequences of proteins obtained from UniProt and submitted to the AlphaFold server for structure prediction and protein-protein interaction analysis. The resulting models were analyzed using PyMOL, and those with the highest confidence scores were selected for further analysis. To identify residues located at the interaction interface between two proteins, the selected models were further processed using InterfaceResidues, an open-source Python script (91). A residue is classified as an interface residue when its accessible solvent-accessible surface area (ΔASA) changes by at least 1.0 Å² after the two chains are separated. The buried surface area (BSA) and binding free energy (ΔG_bind) were calculated using PDB PISA (92). Subsequently, manual calculations in PyMOL were performed to validate the observed trends. These manual calculations were based on the approximate relationship: ΔG_bind ≈ ΔG_solv + ΔG_Hbond + ΔG_salt – TΔS (93). Hydrogen bond and salt bridge parameters obtained from PDB PISA were not incorporated into the PyMOL-based manual calculations due to substantial variability.

### Immunocytochemistry (ICC)

ICC analysis was performed as previously described (35, 36, 43). Briefly, Cultured cells at the desired time points were fixed with 4% paraformaldehyde (PFA) in PBS for 15 min at room temperature (RT) and then permeabilized and blocked for 1 hr in blocking buffer (PBS containing 0.2% Triton X-100 and 3% BSA). They were subsequently incubated overnight with primary antibodies in blocking buffer at 4 °C, followed by washing and incubation with corresponding fluorophore-conjugated secondary antibodies. Cell nuclei were counterstained with Hoechst 33342 (HST, ThermoFisher). Primary antibodies used in this study were listed in Supplemental **Table S3**.

### Fluorescent in situ hybridization (FISH)

FISH was performed as described previously (36, 38, 94). Briefly, cultured cells were washed once with PBS, and fixed with 4% PFA for 30 min at RT. Cells were permeabilized with 0.2% TritonX-100 for 10 min, followed by two washes with PBS. Samples were equilibrated with hybridization buffer composed of 2 X SSC, 10% dextran sulfate, 10 mM ribonucleoside-vanadyl complex (RVC; New England Biolabs) and 20% formamide. A mixture of d*igoxigenin (DIG)-*labeled oligo-dT or dA probes (0.2 ng/µL) and yeast tRNA (0.2 µg/µL) were heated to 95 °C for 5 min and immediately chilled on ice. Probes were then combined with equal volumes of 2 X hybridization buffer and incubated with samples overnight at 37 °C. Anti-DIG antibody (Sigma, 11333089001, 1:100) and corresponding fluorophore-conjugated secondary antibodies (ThermoFisher) were used to detect oligo-dT signals. Anti-MAP2 (Abcam, ab5392, 1:10000) antibody and HST were used to determine neuronal soma and nuclei, respectively. All reagents and solutions were prepared in nuclease-free water.

### Protein nuclear transport assay

Protein nuclear transport was analyzed using a dual reporter (2Gi2R) or individual reporters (RFP-NES, RFP-NLS) as previously described (36, 38, 94). Briefly, cultured cells were transduced with lentiviruses expressing reporters. For SH-SY5Y cells and induced neurons, the reporter lentivirus was transduced when cell growth reached about 50% confluency. The signal density and distribution of GFP and RFP were analyzed at 3 (for SH-SY5Y cells) or 7 (for SH-SY5Y-derived neurons) days post-viral infection (dpi). For iPSC-derived neurons, reporter lentiviruses were co-transduced with lentiviruses expressing MN reprogramming factors. The signal intensity and distributions of GFP and/or RFP were analyzed at desired time points.

### Imaging and quantification analysis

Images were acquired using confocal microscopes (Nikon A1R confocal microscope, and Leica TCS SP5 confocal microscope; 63×/1.4 NA objectives) or an inverted fluorescence microscope (Olympus, CKX53). For co-localization analysis, WT and DYT1 patient fibroblasts were cultured, stained, and imaged under identical confocal settings. Co-localization intensity was quantified based on overlapping foci. Regions of interest were defined by merging the TorsinA (green) and XPO1 (red) channels, followed by application of a color threshold to restrict selection to overlapping (yellow) regions. Correlation coefficients (r) range from −1 to +1, with values closer to ±1 indicating stronger linear relationships and values near 0 indicating weak or no linear association. An r value > 0.7 was considered indicative of a strong positive correlation (95). For quantification of fluorescence intensity of transport reporters, regions encompassing as much of the nucleus or cytoplasm as possible were measured, as previously described (38, 94). For quantification of neurite outgrowth, neurite length was measured using Fiji (ImageJ) software based on TUBB3 staining. The mean branch length was calculated to represent the relative neurite length of each condition. Data collection was performed using an unbiased approach, and image analysis was conducted in a blinded manner.

### Proximity Ligation Assay (PLA)

Cultured cells were fixed with 4% PFA, permeabilized with Triton X-100, and subjected to a proximity ligation assay using antibodies against TorsinA and XPO1. PLA was performed using Duolink In Situ PLA kit according to the manufacturer’s instructions. Nuclei were counterstained with DAPI. PLA puncta were imaged using a Leica TCS SP5 confocal microscope for two-dimensional acquisition and an Olympus spinning disk confocal microscope for three-dimensional imaging. Quantification was performed using Fiji (ImageJ) by counting PLA-positive puncta per cell. Identical thresholding parameters were applied across all experimental conditions to ensure consistency. Negative controls included omission of primary antibodies and single-primary-antibody controls to assess background signals. Data was collected from at least three independent experiments.

### Quantitative real-time PCR analysis

As described previously (36, 43, 96), total RNA was extracted from cultured neurons using TRIzol (Life Technologies). An indirect coculture system was used to prepare highly pure neurons at late maturation stages (>10 dpi) (97). cDNA synthesis reactions were performed using 0.5 µg of total RNA from each sample with the SuperScriptIII First-Strand kit (Life Technologies) and random hexamer primers. Real-time PCR was performed in triplicate using primers, SYBR GreenER SuperMix (Invitrogen), and the BIO-RAD CFX-96 Fast Real-Time PCR system. Target mRNA levels were normalized to the reference gene *GAPDH* or *TUBB3* by 2^-ΔΔCt^ method. Sequences of RT-PCR primers were listed in Supplemental **Table S2.**

### Statistical analysis

Statistical analysis was done in GraphPad Prism. The D’Agostino & Pearson omnibus normality test was conducted first to determine if the data are normally distributed. If the data passed the normality test, one-way or two-way ANOVA was used to determine significance. If the data did not pass the normality test, the Kruskal-Wallis test was used to determine significance. Results are expressed as mean ± SD of at least three biological replicates or three independent experiments, and *P* < 0.05 is considered significant.

## Acknowledgement

We thank members of the Ding laboratory (Drs. Masuma Akter and Dr. Masood Sepehrimanesh) for their technical assistance, and Drs. Mary Munson, Eric First, Shile Huang, and Lucy Robinson for helpful discussions. We also acknowledge support from the Louisiana Optical Network Infrastructure (LONI) for access to supercomputers used in AlphaFold modeling, the Mass Spectrometry Core, Virus Production Core, and Research Core Facilities at LSUHS. This work was supported by National Institutes of Health (NIH) National Institute of Neurological Disorders and Stroke (NINDS) (NS133252 to B.D.), and LSUHS Ike Muslow Predoctoral Fellowship (to H.C).

## Author Contributions

B.D., and H.C. conceptualized and designed experiments; H.C., Y.D., M.K.I., M.A.H., J.L., and X.L. performed experiments; B.D., H.C., Y.D. analyzed and interpreted data; B.D. wrote the original manuscript; B.D. and H.C. edited the manuscript. All authors have read and provided inputs to the manuscript.

