## Supplemental information for "Dystonia-associated TorsinA-ΔE mutation induces a gain-of-function interaction with XPO1 via its N-Terminal hydrophobic segment"


**This PDF file includes:**
Supplementary text
Figures S1-S7
Table S1-S7
References for SI reference citations

Note: Tables S6 and S7 were submitted separately as Excel files.

### Supplementary Information Text

#### Cell lines and culture condition

HEK 293T cells (CRL-11268) and SH-SY5Y (CRL-11266) cells were purchased from ATCC. DYT1 fibroblast cell lines and sex- and age-matched healthy controls were obtained from the *Coriell Institute for Medical Research* or *NINDS Human Cell and Data Repository* (NHCDR). Human induced pluripotent stem cell (hiPSC) lines have been generated in our previous studies (1-3). All hiPSCs were maintained in complete mTeSR1 medium (STEMCELL Technologies) on Matrigel (Corning) coated dishes at 37 °C and 5% $CO_2$ with saturating humanity, and the medium was daily replaced.

HEK medium: DMEM (Gibco) supplemented with 10% fetal bovine serum (FBS, Corning, NY, USA) and 1% penicillin/streptomycin (ThermoFisher).

Fibroblast medium: DMEM supplemented with 15% FBS and 1% penicillin/streptomycin.

hiPSC medium: complete mTeSR1 medium and 1% penicillin/streptomycin.

KOSR medium: DMEM/F12 medium with 20% KnockOut Serum Replacement (KOSR, Thermo Fisher Scientific, MA, USA), 1% GlutaMax, 1% non-essential amino acid (NEAA), 50 μM β-mercaptoethanol (β-ME), 1% P/S and 10 ng/ml basic fibroblast growth factor (bFGF, PeproTech, NJ, USA).

Neurosphere medium (NSP medium): DMEM/F12 medium containing 1% N2, 1% GlutaMax, 1% NEAA, 50 μM β-ME, 1% P/S, 8 μg/ml Heparin, 20 ng/ml bFGF and 20 ng/ml epidermal growth factor (EGF, PeproTech).

Neural progenitor cell medium: DMEM/F12 and neurobasal medium (1:1) containing 0.5% N2, 1% B27, 1% GlutaMax, 1% NEAA, 50 μM β-ME, 1% penicillin/streptomycin, 10 ng/mL EGF and 10 ng/mL bFGF.

Neuronal maturation medium: DMEM: F12: neurobasal (2:2:1), 1% N2 (Invitrogen), 1% B27 (Invitrogen), 1% penicillin/streptomycin, and supplemented with 5 mM FSK and 10 ng/mL each of BDNF, GDNF, and NT3 (PeproTech).

SH-SY5Y medium: DMEM supplemented with 15% FBS and 1% penicillin/streptomycin.

SH-SY5Y differentiation medium: DMEM supplemented with 3% FBS, 10 μM all-trans-Retinoic acid (Sigma) and 1% penicillin/streptomycin.

#### Plasmid construction and lentivirus production

A third-generation lentiviral vector (*pCSC-SP-PW-IRES-GFP*) was used to express reprogramming factors NGN2-IRES-ISL1-T2A-LHX3 (Addgene #221814) as previously reports (4, 5). The same lentiviral vector was used to express GFP-tagged TorsinA, TorsinA variants, and hydrophobic peptides. PCR amplified GFP sequence with incorporated BamHI (5') and NheI (3') sites, and amplified cDNA sequences with NheI (5') and XhoI (3') restriction sites. Inserts of GFP and gene of interest were ligated into the lentiviral vector between the BamHI and XhoI restriction sites. Another third-generation lentiviral vector (*LV-CAG-mCherry-miRE-Luc*) was used to express shRNAs targeting TOR1AIP1 (LAP1) and TOR1AIP2 (LULL1) as described in previously reports (2, 4, 6). Briefly, shRNA oligos targeting the gene coding regions were synthesized, PCR amplified with flanking miR-30 sequences, and ligated into the above lentiviral vector at the *XhoI* and *EcoRI* restriction sites. The targeting sequences (gRNAs) were listed in Supplemental **Table S2**. The dual reporter 2Gi2R (Addgene #71396) was generously provided by Dr. Fred H. Gage (7). The transport reporters of RFP-NES (Addgene #221812) and RFP-NLS (Addgene #221813) were constructed as previously reported (4).

All plasmids were verified by restriction enzyme digestions and DNA sequencing. Each of these vectors was co-transfected with packaging plasmids (Addgene #12251, #12253, and #12259) into HEK293T cells for virus production (8, 9). Replication-incompetent lentiviruses were produced, and viral supernatants were collected at 48 hrs and 72 hrs post-transfection as previously described (10, 11). The viral supernatants were filtered through 0.45 μm syringe filters and stored at 4 °C prior to cell transduction.

**Supplementary Figures and Figure Legends**

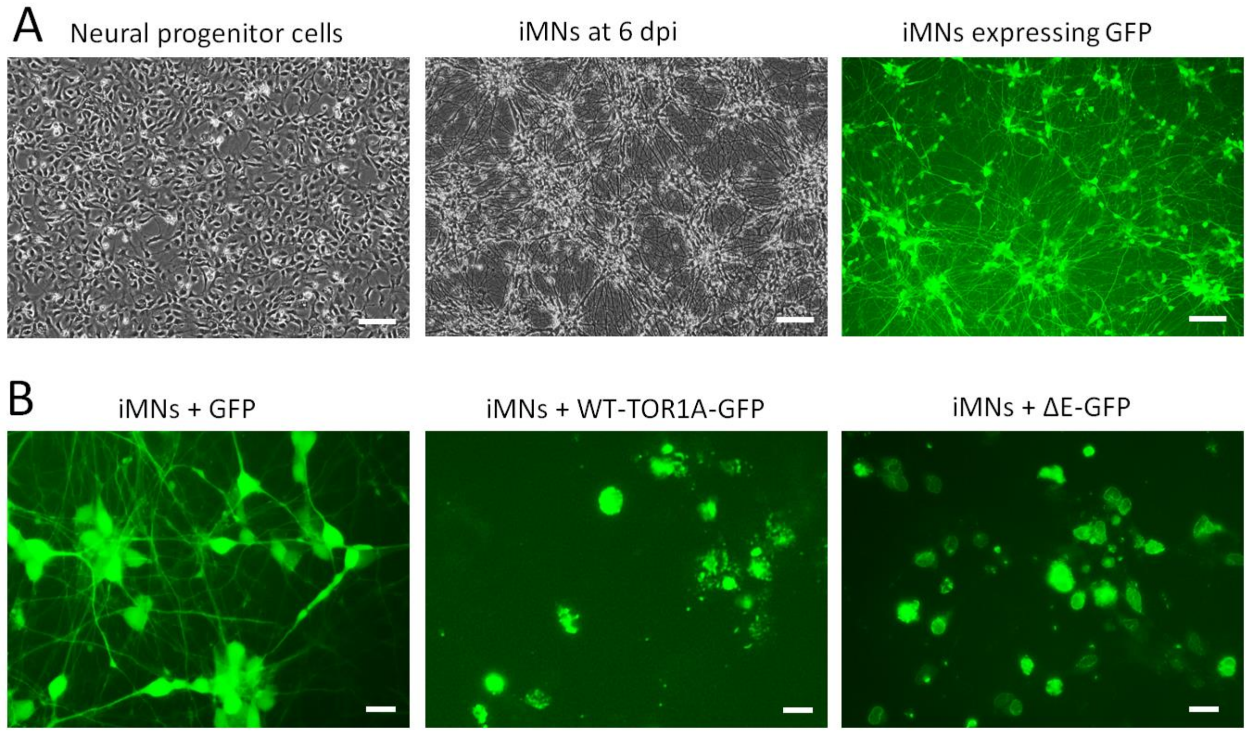


**Figure S1. Generation of induced human motor neurons expressing GFP-tagged WT and mutant TorsinA (ΔE).**

**(A)** Representative low-magnification images of neural progenitor cells (NPCs) at the time of lentiviral transduction, induced motor neurons (iMNs) at 6 days post-viral infection (dpi), and GFP expression in iMNs at 6 dpi. Scale bars, 100 µm.

**(B)** Representative high-magnification fluorescence images of iMNs expressing GFP, WT TorsinA-GFP, or TorsinA-ΔE-GFP at 6 dpi. Scale bars, 20 µm.

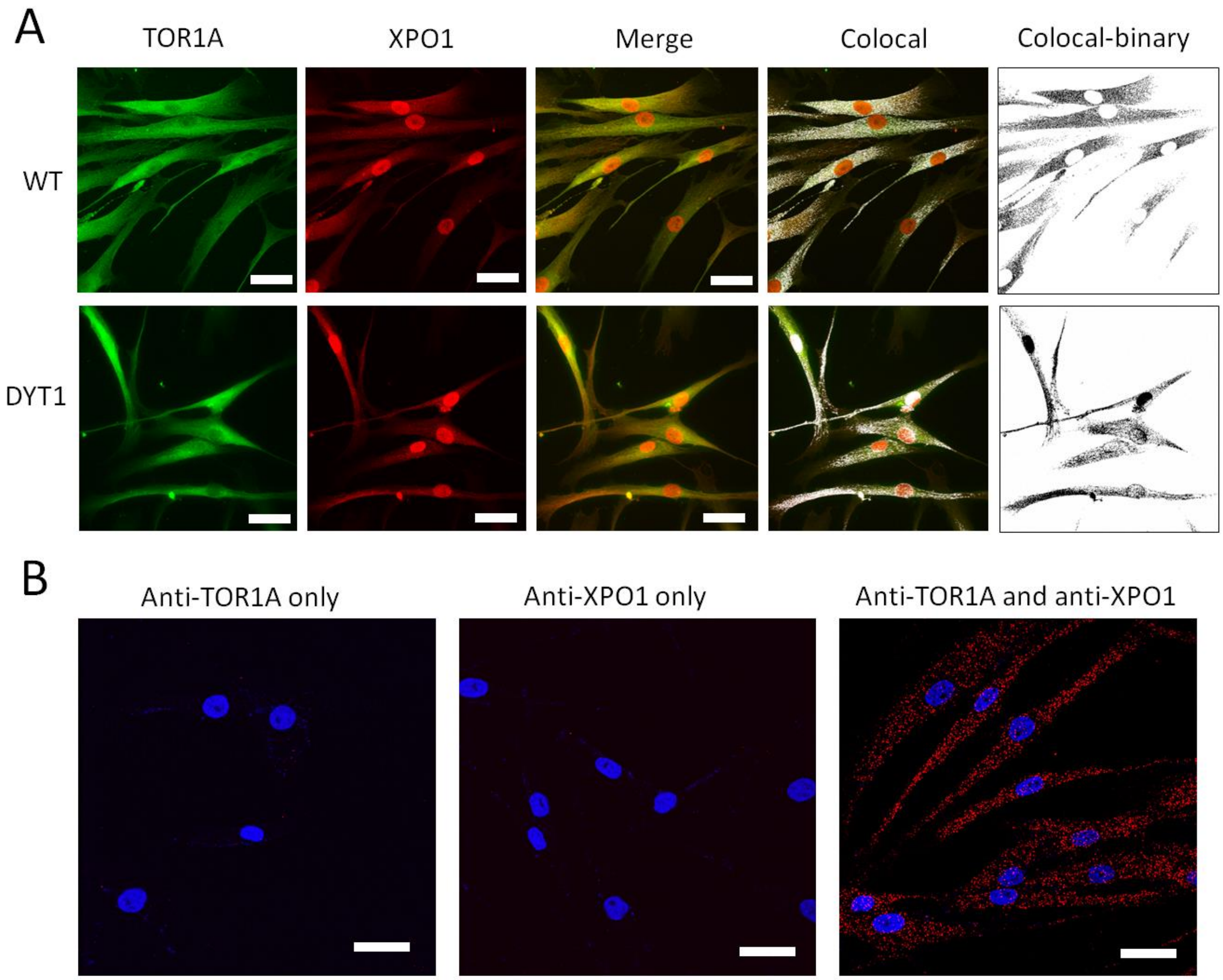


**Figure S2. Colocalization and proximity of endogenous TorsinA and XPO1 in human fibroblasts.**

(**A**) Representative low-magnification immunofluorescence images showing the subcellular localization and colocalization of endogenous TorsinA and XPO1 in control (WT; GM04506) and DYT1 patient-derived (GM03211) fibroblasts. Scale bars, 20 µm.

(**B**) Negative-control proximity ligation assays (PLA) performed in WT fibroblasts with either the TorsinA or XPO1 primary antibody omitted to evaluate assay specificity. Scale bars, 20 µm.

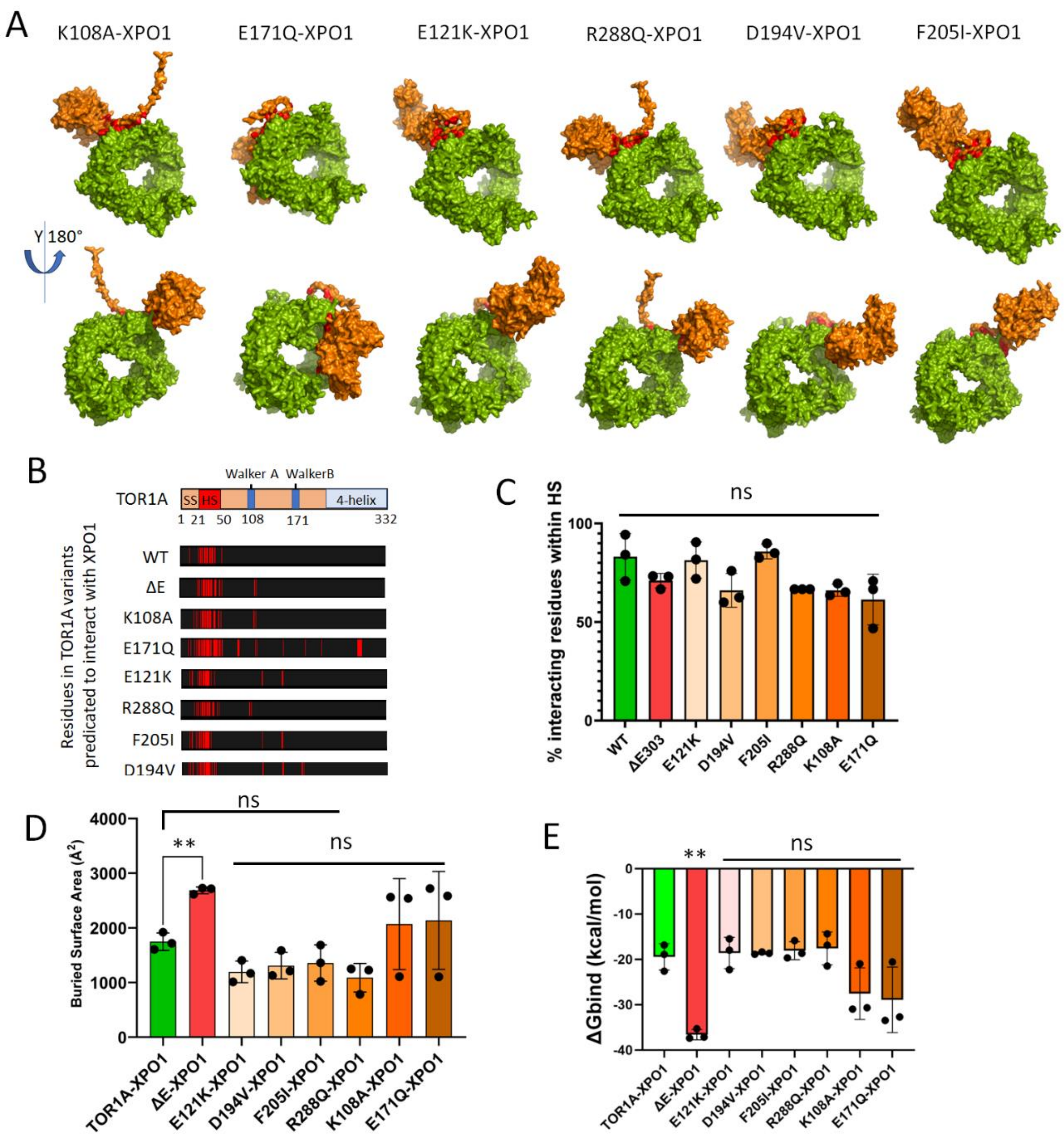


**Figure S3. Structural modeling predicts that enhanced XPO1 association is specific to TorsinA-ΔE among the variants examined.**

(**A**) Highest-confidence AlphaFold-predicted structural models of the indicated TorsinA variants in complex with XPO1.

(**B**) Predicted XPO1 interface residues in each TorsinA variant, highlighted in red, from the highest-confidence model of each complex.

(**C**) Percentage of predicted TorsinA interface residues located within the N-terminal HS for each TorsinA-XPO1 complex. Values were derived from the three highest-confidence models generated for each variant.

(**D**) Predicted buried surface areas of the indicated TorsinA-XPO1 complexes, calculated from the three highest-confidence models.

(**E**) Estimated binding free energies ($\Delta G\_bind$) of the indicated TorsinA-XPO1 complexes, calculated from the three highest-confidence models.

For (**C-E**), compared to WT TorsinA-XPO1, ns, not significant; ** $p<0.01$. One-way ANOVA with Dunnett's multiple comparison test.

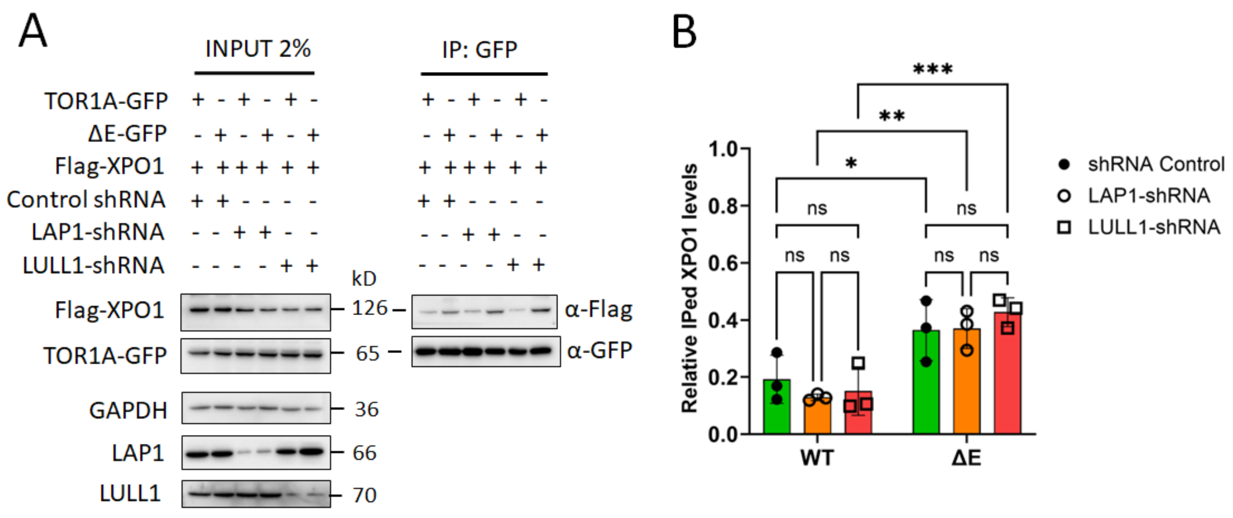


**Figure S4. Depletion of the TorsinA cofactors LAP1 or LULL1 does not alter TorsinA-XPO1 association.**

**(A)** Co-immunoprecipitation analysis of GFP-tagged WT TorsinA and TorsinA-ΔE in HEK cells coexpressing FLAG-tagged XPO1 and shRNAs targeting the TorsinA cofactors LAP1 or LULL1. GFP-tagged TorsinA proteins were immunoprecipitated using GFP-Trap and analyzed by immunoblotting with the indicated antibodies.

**(B)** Quantification of co-immunoprecipitated FLAG-XPO1 normalized to the corresponding GFP-tagged TorsinA bait. Data are from three independent experiments. ns, not significant; $*P < 0.05$, $**P < 0.01$, and $***P < 0.001$; one-way ANOVA followed by Dunnett's multiple-comparisons test.

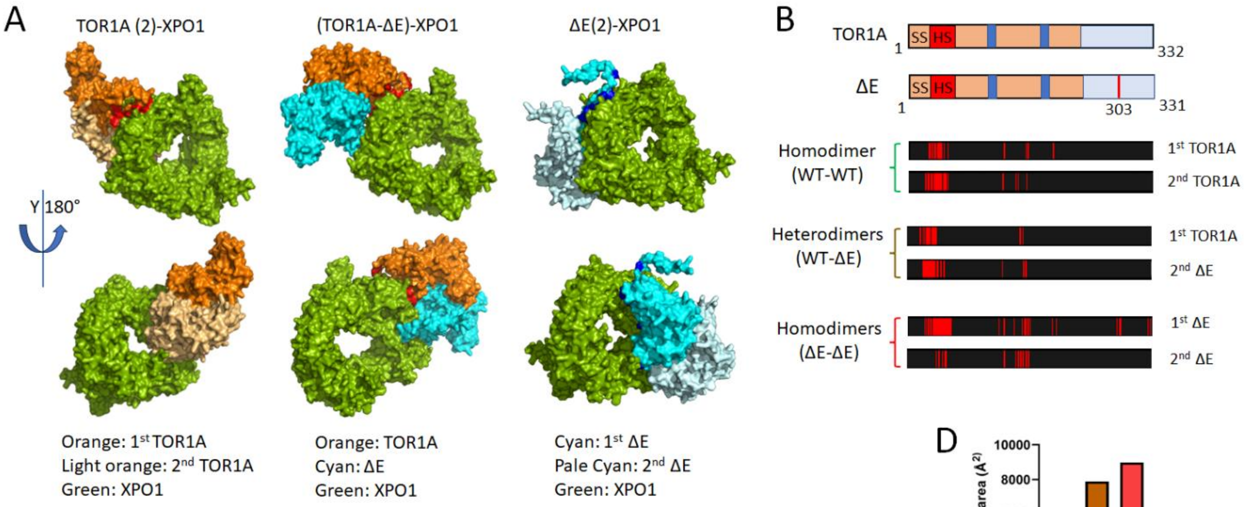


C

| WT homodimer: TOR1A(2)-XPO1 | | | | |
|---|---|---|---|---|
| 1st TOR1A | 2nd TOR1A | 1st AA # | 2nd AA # | BSA (Å²) |
| VEPISLGLALAGVLTGYIYPRLYC<br>LFAECC--DY--GNVSA--GV | VEPISLGLALAGVLTGYIYPRLYCL<br>FAECCGQK–DY–LYKD--SA | 25 | 33 | 6264 |
| 21-50; 127-128; 157-161;<br>195-196 | 21-53;217-128; 146-149;<br>160-161 | | | |
| **Heterodimer: (TOR1A-ΔE)-XPO1** | | | | |
| 1st TOR1A | 2nd ΔE | 1st AA # | 2nd AA # | BSA (Å²) |
| VEPISLGLALAGVLTGYIYPRLYC<br>LFA--LWIRG | VEPISLGLALAGVLTGYIYPRLYCL<br>FA--D--RGNVSA | 20 | 27 | 7904 |
| 21-47; 153-157 | 21-47; 127; 156-161 | | | |
| **ΔE homodimer: ΔE(2)-XPO1** | | | | |
| 1st ΔE | 2nd ΔE | 1st AA # | 2nd AA # | BSA (Å²) |
| VEPISLGLALAGVLTGYIYPRLYC<br>LFAECCGQKRS--E-DY-S-<br>LWIRGNVSACAR--D-K--<br>EMQSRG--D-D | LTGYIYPRLYCLFAE--DY--<br>NITLYKDQLQLWIRGNVSA | 50 | 21 | 8994 |
| 21-55; 121; 127-128;<br>140;153-164; 194; 200;<br>284-289; 326; 330 | 34-48; 127-128; 143-161 | | | |

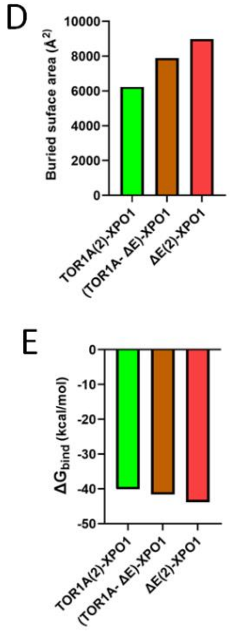


**Figure S5. The enhanced interaction of ΔE with XPO1 within dimers.**

(**A**) AlphaFold models of interactions between TOR1A and/or ΔE dimers and XPO1, including WT homodimer TOR1A(x2), WT/ΔE heterodimer TOR1A-ΔE, and ΔE homodimer ΔE(x2).

(**B**) Predicated interacting residues (red) in TOR1A variants when associated with XPO1 as dimers.

(**C**) Summary of protein-protein interacting residues in TOR1A variants when associate with XPO1 as dimers based on models with the highest confident scores.

(**D**) The buried surface area of indicated complexes. Values represent the models with the highest confident scores.

(**E**) The change in Gibbs free energy of binding ($\Delta G_{bind}$) in indicated complexes based on models with the highest confident scores.

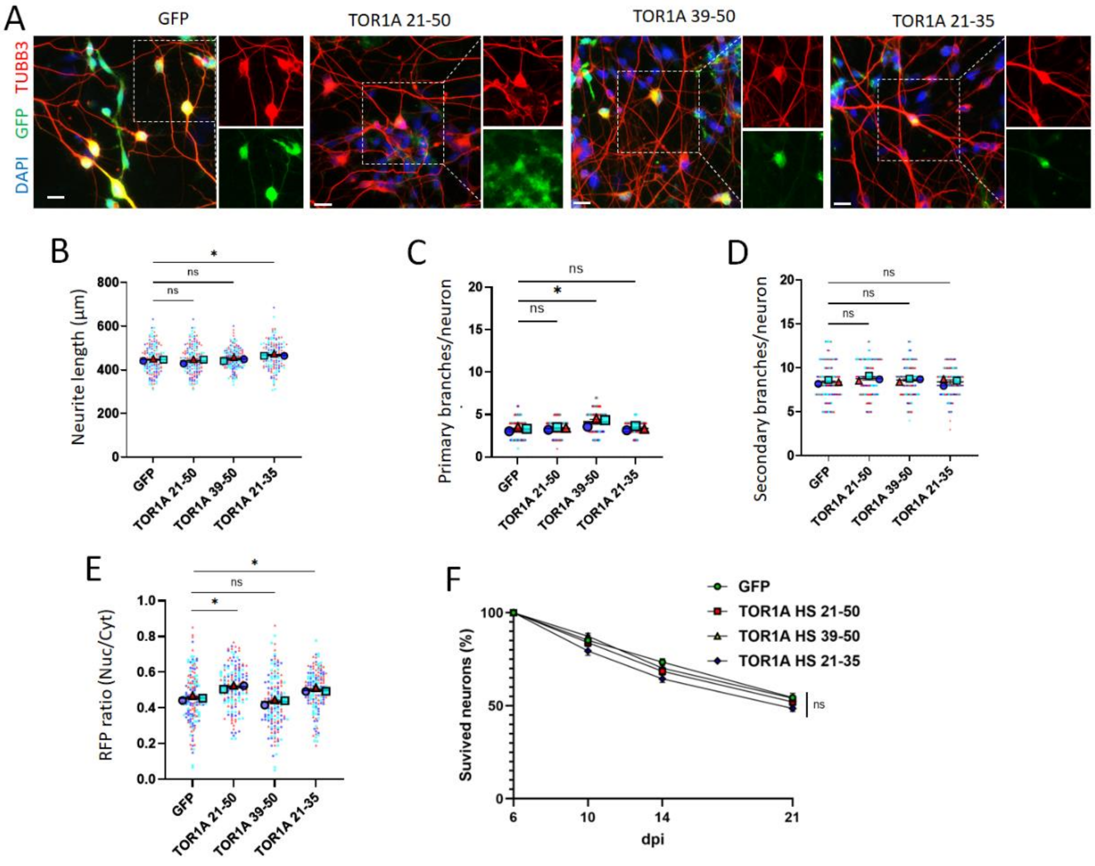


**Figure S6. Expression of TorsinA HS-derived peptides produces modest effects in healthy control motor neurons.**

(**A**) Representative fluorescence images of control hiPSC-derived motor neurons transduced with lentiviral vectors expressing GFP or the indicated GFP-tagged TorsinA HS-derived peptides. Motor neurons were cocultured with astrocytes and imaged at 6 days post-induction (dpi). Scale bars, 20 μm.

(**B-D**) Quantification of (**B**) neurite length at 6 dpi, (**C**) number of primary branches at 10 dpi, and (**D**) number of secondary branches at 10 dpi in control hiPSC-derived motor neurons expressing GFP or the indicated HS-derived peptides.

(**E**) Quantification of nuclear export using the RFP-NES reporter in control hiPSC-derived motor neurons at 6 dpi. Nuclear export was assessed by measuring the nuclear-to-cytoplasmic ratio of RFP-NES fluorescence.

For (**B-E**), 150 neurons per group were analyzed across three independent experiments, with data from each experiment shown in a distinct color. ns, not significant; **P* < 0.05; one-way ANOVA followed by Dunnett's multiple-comparisons test.

(**F**) Survival of control hiPSC-derived motor neurons expressing GFP or the indicated GFP-tagged HS-derived peptides. The number of surviving neurons at 6 dpi was defined as 100% for each condition, and subsequent measurements were normalized to this baseline. A total of 200 neurons per group were analyzed across three independent experiments. ns, not significant; one-way ANOVA followed by Dunnett's multiple-comparisons test.

### Supplementary Tables

#### Table S1. List of cell lines used in this study and their Research Resource Identifiers (RRIDs).

| | Resource | Sex | Age at Sampling (YR) | Tissue Type | Race | Onset Age (YR) | Gene | Mutation |
|---|---|---|---|---|---|---|---|---|
| **Fibroblast cell lines** | | | | | | | | |
| DYT1-1 | Coriell Cat# GM03211, RRID:CVCL_1U24 | M | 30 | Skin | Caucasian | 7 | TOR1A | 907_909delGAG |
| DYT1-2 | NDS00305 | M | 22 | Skin | Caucasian | 12 | TOR1A | 907_909delGAG |
| DYT1-3 | Coriell Cat# GM02304, RRID:CVCL_1U21 | F | 17 | Skin | Caucasian | 7 | TOR1A | 907_909delGAG |
| DYT1-4 | NDS00301 | F | 53 | Skin | Caucasian | 9 | TOR1A | 907_909delGAG |
| Control-1 | Coriell Cat# GM00024, RRID:CVCL_7269 | M | 31 | Skin | Caucasian | | | |
| Control -2 | Coriell Cat# GM03652, RRID:CVCL_7397 | M | 24 | Skin | Caucasian | | | |
| Control -3 | Coriell Cat# GM04506, RRID:CVCL_7413 | F | 20 | Skin | Caucasian | | | |
| Control -4 | Coriell Cat# AG07473, RRID:CVCL_2C33 | F | 50 | Skin | Caucasian | | | |
| **hiPSC lines** | | | | | | | | |
| DYT1-1 | DYT1-1C8, Ref. (3) | M | 30 | TOR1A 907_909delGAG | | | | |
| DYT1-2 | DYT1-H6, Ref. (3) | M | 30 | TOR1A 907_909delGAG | | | | |
| WT-1 | WT-1A2, Ref. (3) | M | 30 | Isogenic control for 1C8 and H6 | | | | |
| WT-2 | WT-1F6, Ref. (3) | M | 30 | Isogenic control for 1C8 and H6 | | | | |
| DYT1-3 | DYT1-11, Ref. (2) | M | 30 | TOR1A 907_909delGAG | | | | |
| WT-3 | WT-4B2, Ref. (1) | M | 30 | Isogenic control for DYT1-11 | | | | |
| WT-4 | WT-A6, Ref. (1) | M | 30 | Isogenic control for DYT1-11 | | | | |
| **HEK 293T cells** | | ATCC Cat# CRL-11268, RRID:CVCL_1926 | | | | | | |
| **SH-SY5Y cells** | | ATCC, Cat# CRL-11266, RRID:CVCL_0019 | | | | | | |

Note: RRIDs, Research Resource Identifiers (RRIDs); M, Male; F, Female.

### Table S2. Oligo Sequence

| Name | Sequence (5’-3’) | Note |
|---|---|---|
| TOR1AIP1-shRNA-1 | CAGCAGTGTCACTACTGTTAA | gRNA |
| TOR1AIP1-shRNA-2 | CAAGACAGTGATACTGTCAAA | gRNA |
| TOR1AIP2-shRNA-1 | ACCATGAGACAGATTAATAAA | gRNA |
| TOR1AIP2-shRNA-2 | CGGCTCCACTTTGATCTTCTATA | gRNA |
| 5'TOR1A-BamHI | ggatccGCCACCATGAAGCTGGGCCGGGC | Truncate Cloning |
| 3'TOR1A311-NheI | GTTgctagc TCCTCTTTGGGGAAAAATGT | Truncate Cloning |
| 3'TOR1A250-NheI | GTTgctagc CTGTTCTTGTTATTGAAAACCG | Truncate Cloning |
| 3'TOR1A90-NheI | GTTgctagc TGGGTTGTTTATGAAACCAAA | Truncate Cloning |
| 5’TOR1A91-BamHI | Gtt ggatcc GCCACC ATG AAGCCCAAGAAACCTCTC | Truncate Cloning |
| 3'TOR1A-NheI | GTTgctagc ATCATCGTAGTAATAATC | Truncate Cloning |
| 5’TOR1A51-BamHI | ggatccGCCACCatgGGGCAGAAGCGGAGCCTTAG | Truncate Cloning |
| 5’TOR1A21-BamHI | ggatccGCCACCatgGTGGAGCCCATCAGCCTGG | Truncate Cloning |
| 5’TOR1AΔ21-50 | TCTGCCCCGCCTGCACCACGGACGG | Truncate Cloning |
| 3’TOR1AΔ21-50 | TGCAGGCGGGGCAGAAGCGGAGCCTTAGC | Truncate Cloning |
| 5’TOR1A 21-BamHI | GGATCCGCCACCATGGTGGAGCCCATCAGCCTG | Peptide cloning |
| 5’TOR1A 39-BamHI | GGATCCGCCACCATGTACCCGCGTCTCTACTGCC | Peptide cloning |
| 3’TOR1A 50-NheI | GTT GCTAGCGCAGCACTCGGCGAAGAGG | Peptide cloning |
| 3’TOR1A 48-NheI | GTT GCTAGCCTCGGCGAAGAGGCAGTAG | Peptide cloning |
| 3’TOR1A 35-NheI | GTT GCTAGCGGTGAGGACGCCGGCCAG | Peptide cloning |
| CHAT-F | ACAACCACGGAGATGTTCTG | RT-PCR |
| CHAT-R | TGCAGCTGTGAAAGCTAGAG | RT-PCR |
| GAPDH-F | CAAATTCCATGGCACCGTCA | RT-PCR |
| GAPDH-R | GGACTCCACGACGTACTCAG | RT-PCR |
| MAP2-F | ACTCCTGGAACCCCTAGCTA | RT-PCR |
| MAP2-R | TGGGAGTCGCAGGAGATTTT | RT-PCR |
| NEFH-F | TGAGCTGAGGAACACCAAGT | RT-PCR |
| NEFH-R | AGCCAATCCGACACTCTTCA | RT-PCR |
| RBFOX3 (NEUN)-F | CTTACGGAGCGGTCGTGTAT | RT-PCR |
| RBFOX3 (NEUN)-F | TCACATGGTTCCAATGCTGT | RT-PCR |
| SYN1-F | AATACTGGCTCTGCGATGCT | RT-PCR |
| SYN1-R | TGACCACGAGCTCTACGATG | RT-PCR |
| Oligo-dT | TTTTTTTTTTTTTTTTTTTTT-DIG | FISH |
| Oligo-dA | AAAAAAAAAAAAAAAAAAAAA-DIG | FISH |

Note: DIG, Digoxigenin; FISH, Fluorescence *In Situ* Hybridization.

**Table S3. List of primary antibodies used this study.**

| Name | Source | Catalogue | Host | Immunostaining | Western Blot |
|---|---|---|---|---|---|
| β-Actin | Sigma | A5441 | Ms | - | 1:5000 |
| DIG | Sigma | 11 333 089 001 | Sh | 1:100 | - |
| FLAG | Sigma | F1804 | MS | - | 1:3000 |
| GAPDH | Proteintech | 50-553-944 | MS | - | 1:2000 |
| GFP | Aves | GFP-1020 | Ck | 1:1000 | 1:5000 |
| MAP2 | Abcam | Ab5392 | Ck | 1:10000 | - |
| TUBB3 | Covance | MMS-435P | Ms | 1:2000 | - |
| TUBB3 | Covance | PRB-435P | Rb | 1:2000 | - |
| TOR1A | Cell Signaling | 2150 | Ms | 1:200 | 1:1000 |
| TOR1AIP1 | Sigma | 50546 | Rb | - | 1:500 |
| TOR1AIP2 | Protein tech | 24769-1-AP | Rb | - | 1:300 |
| XPO1 | Sigma | HPA042933 | Rb | 1:200 | 1:1000 |

Note: Ck, chicken; Gt, goat; Ms, mouse; Rb, rabbit; Sh, sheep; - not determined in this study.

**Table S4. AlphaFold predicated residues in TOR1A or ΔE when interacting with indicated proteins.**

| | | | % interacting residues within HS | |
|---|---|---|---|---|
| | **WT-TOR1A** | **ΔE** | **In TOR1A** | **In ΔE** |
| **XPO1** | VEPISLGLALAGVLTGYIYPRLYCL FAECCGQKRS | VEPISLGLALAGVLTGYIYPRLYCLFA ECCGQKRSLSREALQKD--ENIYE | **94.4** | **66.7** |
| | 21-50 | 21-64; 117-121 | | |
| **CANX** | VEPISLGLALAGVLTGYIYPRLYCL FAECC | VEPISLGLALAGVLTGYIYPRLYCLFA ECC | **32** | **35.1** |
| | 21-50 | 21-50 | | |
| | EGGLNSDYVHLFVATLHFPHASN ITLYKDQLQLWIRGNVSA | LFVATLHFPHASNITLYKDQLQLWI RGNVSA | | |
| | 121-161 | 131-161 | | |
| | PLEYKHLKMCIRVEMQSRGYEID EDIVSRVAEEMT | PLEYKHLKMCIRVEMQSRGYEIDE DIVSRVA | | |
| | 271-305 | 271-301 | | |
| **CLGN** | VEPISLGLALAGVLTGYIYPRLYCL FAECC | VEPISLGLALAGVLTGYIYPRLYCLFA ECC | **26** | **27.1** |
| | 21-50 | 21-50 | | |
| | EGGLNSDYVHLFVATLHFPHASN ITLYKDQLQLWIR | LYKDQLQLWIR | | |
| | 121-156 | 146-156 | | |
| | FWRSGKQREDIKLKDIEHALSVS VFNNKNSGFWHSSLIDRN | DAIKPFLDYYDLVDGVSYQKA-- DFWRSGK | | |
| | 221-261 | 181-201; 220-226 | | |
| | AEEMTFFPKEERVFSDKGCKTVF TKLDYYY | YKHLK--- DIVSRVAEMTFFPKEERVFSDKGCK TVFTKLDYYYDD | | |
| | 301-330 | 274-278; 295-331 | | |
| **UNC13B** | HASNI | HLFVATLHFPHASNITLYKDQLQLW | **0** | **0** |
| | 140-144 | 132-156 | | |
| | DFWRSGKQREDIKLKDIEHALSV SVFNNKNSGFWHSSLIDRN | GAERITDVALDFWRSGKQREDIKLK DIEHALSVSVFNNKNSG | | |
| | 220-261 | 210-251 | | |
| | EEMTFFPKEER | | | |
| | 302-312 | | | |

Note: Based on models with the highest confidence scores. Predicated interacting residues are highlighted in red.

**Table S5. AlphaFold predicated residues in TosinA variants when interacting with XPO1.**

| TorsinA variants | Consequences and disease | PPI sequences and interacting residues (in red) | Total PPI residues | PPI residues within HS | % residues within HS |
|---|---|---|---|---|---|
| **WT** | N/A | VEPISLGLALAGVLTGYIYPRLYCL FAECCGQKRS<br>21-55 | 18 | 17 | 94.4 |
| **ΔE303** | Loss-of-function; Childhood-onset generalized DYT1 dystonia | VEPISLGLALAGVLTGYIYPRLYCL FAECCGQKRSLSREALQKD-- ENIYE<br>21-64; 117-121 | 24 | 16 | 66.7 |
| **K108A** | Fail to bind ATP | VEPISLGLALAGVLTGYIYPRLYCL FAECCGQKRSLSREALQKD-- ENIYE<br>21-64; 117-121 | 23 | 15 | 65.2 |
| **E171Q** | Fail to hydrolyze ATP | VEPISLGLALAGVLTGYIYPRLYCL FAECCGQKRSLSREALQKD-- NPK--NIYE--R--K--Q--QSRGYEI<br>21-64; 89-91; 118-121; 164; 200; 227; 286-292 | 47 | 22 | 46.8 |
| **E121K** | Reduced TOR1A stability; focal dystonia | VEPISLGLALAGVLTGYIYPRLYCL FAECC--DY--SA<br>21-50; 127-128; 160-161 | 22 | 18 | 81.8 |
| **R288Q** | Similar to ΔE; childhood-onset generalized dystonia | VEPISLGLALAGVLTGYIYPRLYCL FAECCQKRSLSREALQKD--ENIYE<br>21-64; 117-121 | 24 | 16 | 66.7 |
| **F205I** | Disrupt interaction with cofactors and oligomerization; sporadic late-onset dystonia. | VEPISLGLALAGVLTGYIYPRLYCL FAECC--Y--SA<br>21-50; 128; 160-161 | 20 | 17 | 85 |
| **D194V** | Impairs interaction with cofactor LAP1; Childhood-onset dystonia. | VEPISLGLALAGVLTGYIYPRLYCL FAECC--DY--SA--LVVG<br>21-50; 127-128; 160-161; 192-195 | 25 | 19 | 76 |

Note: Based on models with the highest confidence scores. Predicated interacting residues are highlighted in red.

**Table S6. List of proteins identified by Co-IP/MS**
Submitted separately as an Excel file.

**Table S7. Detailed Gene Ontology (GO) analysis results of identified proteins**
Submitted separately as an Excel file.
